# Intertwined degradation of plant diversity and permafrost in sub-Arctic palsa mires

**DOI:** 10.64898/2026.09.07.749784

**Authors:** Oona Leppiniemi, Mika Jokikokko, Julia Kemppinen, Tuija Maliniemi

## Abstract

1. Plant communities of sub-Arctic and Arctic regions are undergoing changes due to rapid climate change. Permafrost degradation is one of the key drivers of habitat change in these environments and it is considered to impact the biodiversity of northern peatlands. However, the empirical evidence linking permafrost thaw to biodiversity change remains limited.
2. Here, we conducted a vegetation survey of the plant communities (including vascular plants, bryophytes, and lichens) of 14 palsa mires to quantify biodiversity change along different stages of permafrost degradation. We quantified changes in community composition, indicator species, alpha- and beta diversity as well as in two biodiversity indicators.
3. Our results showed that permafrost degradation significantly affected both the plant community composition and the biodiversity of palsa mires. The most pronounced changes in biodiversity were associated with mires from which the permafrost had thawed completely. However, those mires that represented high stage of degradation but still retained remnant permafrost features, were still maintaining plant communities comparable to palsas with low stages of degradation. Thus, our results indicate that collapsing palsas are a clear tipping point for local biodiversity. Almost a complete loss of lichens was observed in completely thawed mires. Furthermore, the shifts in the vascular plant community were recognized to have potential to affect the biodiversity of palsa mires beyond vegetation.
4. **Synthesis:** This study underscores the role of permafrost sustaining the compositional and taxonomic diversity of plants in palsa mires while also highlighting the varying responses of vascular plants, bryophytes, and lichens to permafrost degradation. Overall, our findings illustrate how the climate-induced changes in habitats and loss of environmental heterogeneity impact the biodiversity and the ecological resilience of the permafrost peatlands.

## 1. Introduction

The biodiversity of Arctic peatlands remains poorly quantified, even though these high-latitude ecosystems are warming two to four times faster than the global mean temperature (Rantanen et al., 2022). This is a significant knowledge gap because climate-driven permafrost degradation threatens the distinct habitats provided by Arctic peatlands, affecting both their biodiversity and critical ecosystem processes such as greenhouse gas emissions (Cox et al., 2025; Errington et al., 2024; Miner et al., 2022). Palsa mires are considered the most climate-sensitive permafrost peatland ecosystems as they occur in the southern margin of the circumpolar permafrost region (Leppiniemi et al., 2023; Seppälä, 1988). Palsa mires and other major periglacial processes are predicted to be almost completely lost by 2100 across northern Europe due to climate warming (Aalto et al., 2014, 2017; Fewster et al., 2022). Furthermore, the suitable environments for palsa mires are projected to disappear even at the circumpolar scale (Leppiniemi et al., 2023), and the rapid permafrost degradation has led to the classification of palsa mires as critically endangered habitats in Europe (Janssen et al., 2016).

Palsa mires are characterized by large permafrost-cored mounds referred to as ‘palsas’ or ‘peat plateaus’ depending on their morphology (Åhlman, 1977). These permafrost features create microtopographical and hydrological variation within the mire, resulting in sharp environmental gradients and several different microhabitats between hummock and flark-levels (Beilman, 2001; Camill, 1999; Luoto et al., 2004). Such fine-scaled heterogeneity in the palsa habitats supports biodiversity, including rich birdlife, abundant invertebrates, and distinctive plant communities (e.g., Beilman, 2001; Järvinen & Sammalisto, 1976; Markkula, 2014; Zuidhoff & Kolstrup, 2005). However, quantitative assessments of palsa mire biodiversity remain limited (see Cox et al., 2025; Errington et al., 2024). This gap in knowledge is concerning considering the strong degradation of palsas across the Northern Hemisphere during the past decades (Borge et al., 2017; Mamet et al., 2017; Olvmo et al., 2020; Payette et al., 2004; 2012; Verdonen et al., 2023; Wang et al., 2023). In addition to biodiversity, degradation of palsa mires may also affect Indigenous cultures and livelihoods of locals by fragmenting reindeer winter grazing areas and by reducing the harvest of economically important berries such as cloudberry (*Rubus chamaemorus*) (Wang et al., 2026; Ward Jones et al., 2024).

Vegetation of palsa mires have been studied in different parts of the Northern Hemisphere by using vegetation surveys, aerial imagery, Unoccupied Aerial Vehicles, and analysis of peat profiles and macrofossils (e.g., Ruuhijärvi, 1960; Bhiry & Robert, 2006; Errington et al., 2024; Malmer et al., 2005; Normand et al., 2017; Oksanen, 2008; Wolff et al., 2025). The evidence from these studies shows that palsa mires maintain different plant communities during permafrost aggradation and degradation. As permafrost establishes, vegetation transitions from *Sphagnum*-dominated mire surface communities toward dwarf shrub- and lichen -dominated communities on elevated palsas (Zuidhoff & Kolstrup, 2005). Conversely, degrading palsas exhibit reversed shifts in their community composition, which have also been linked with declines in species richness, as reported in North American palsa mires (Beilman, 2001; Cox et al., 2025; Errington et al., 2024). Beyond these characterizations and observed shifts in plant community composition and species richness, research on the contribution of palsa mires to the different aspects of biodiversity remains limited or is completely lacking in many circumpolar regions.

Given the rapidly changing environmental conditions and the lack of knowledge, more comprehensive quantification of the biodiversity sustained by palsa mires are urgently needed. Biodiversity assessments are increasingly required to consider different aspects of biodiversity to understand the overall outcomes of biodiversity change (Dornelas et al., 2023). Such aspects include e.g. changes in community composition, species identity, dominance, and ecological relevance in addition to species richness that has often been used as the only metric to quantify biodiversity. Relying solely on species richness gives however, a limited view on biodiversity change (Dornelas et al., 2023; Hillebrand et al., 2018; Maliniemi et al., 2025). Without such assessments, the extent of biodiversity loss associated with the permafrost degradation and potential disappearance of palsas, will remain unknown. In this study, we address this gap by quantifying the effect of permafrost thaw on different aspects of plant communities and their biodiversity in palsa mires. By conducting a vegetation survey of 14 palsa mires representing different stages of permafrost degradation in northwestern Finland, Northern Europe, we answer to the following research questions:

1. How do the plant community composition and diversity shift along the distinct stages of permafrost degradation in palsa mires?
2. How do the diversity patterns of different taxonomic groups (i.e. vascular plants, bryophytes, and lichens) vary within and between palsa mires at different stages of permafrost degradation?

Based on the findings of Errington et al. (2024) and Cox et al. (2025), we hypothesize that pristine palsa mires support the most unique plant communities in comparison to more degraded mires, and that this diversity start decline as permafrost thaw advances and the elevated palsas collapse and transition towards wetter, permafrost-free growing conditions, resembling the surrounding fen. Especially, we expect to find a sharp decline in lichen abundance and diversity as permafrost disappears. We therefore anticipate that changes in species composition will reduce plant diversity in palsa mires, illustrating how climate-driven habitat changes can contribute to biodiversity loss.

## 2. Materials and methods

### 2.1. Study area

Our study area locates in the most northwestern part of Finnish Lapland, in Northern Europe, at the southern margin of the permafrost region (Figure 1). We focus on mires that lie within a 60km NW– SW belt stretching from the vicinity of Mount Leutsuvaara (68.92° N, 20.93° E) to lake Kelottijärvi (68.52° N, 21.99° E). The mean annual air temperature in the region is −1.4 °C, and the total annual precipitation is approximately 500 mm (Finnish Meteorological Institute, 2026ab), measured from the nearest meteorological stations (Fig. 1). In general, the environmental conditions are relatively similar across the studied palsa mires, elevation varying from 385–470 m asl. Semi-domesticated reindeer (*Rangifer tarandus tarandus*) is the main large herbivore in the area, where their numbers have been high over several decades (Bernes et al., 2015).

**Figure 1.**
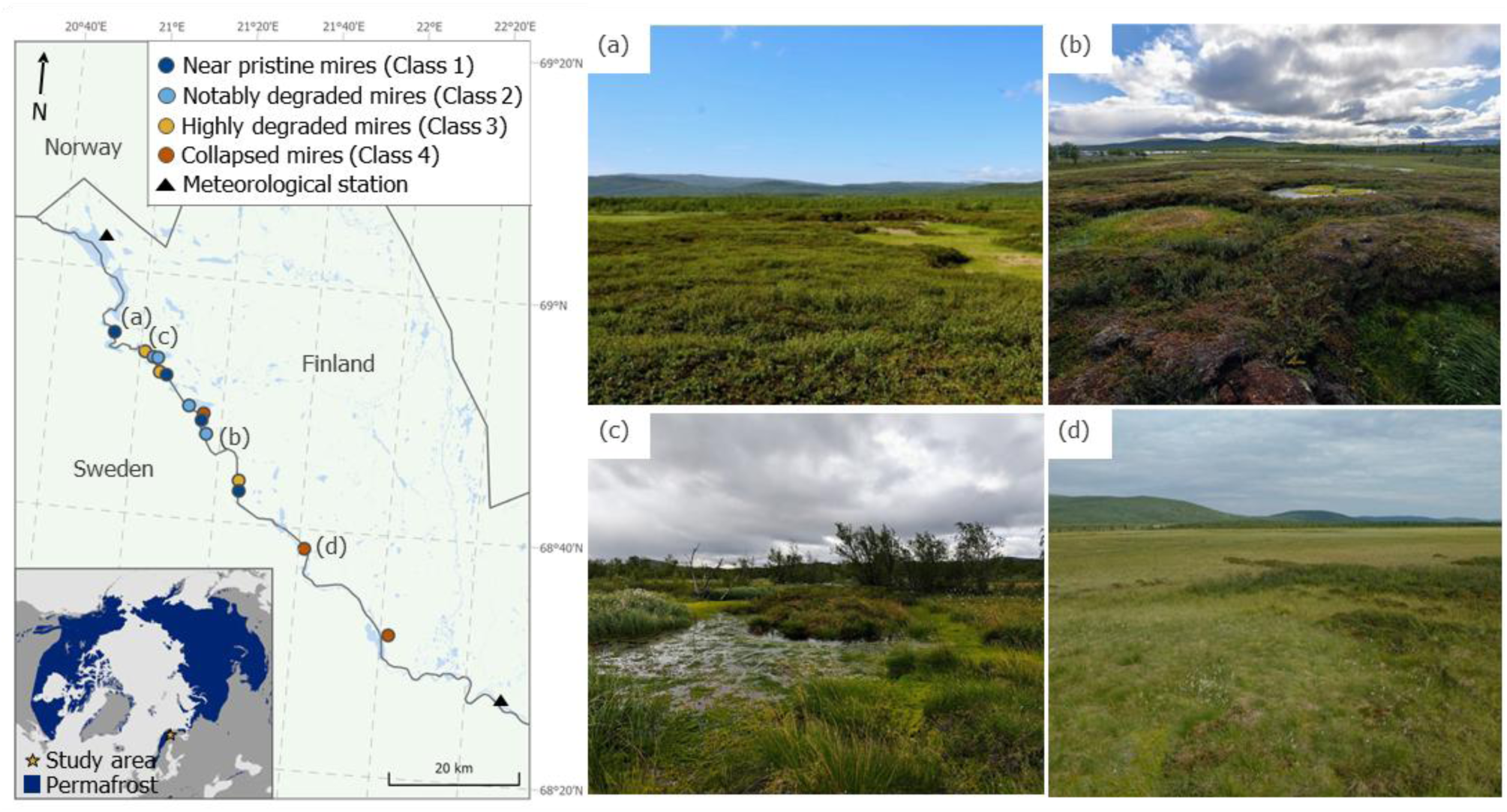
The study area and the locations of the studied palsa mires, classified according to their state of permafrost degradation, are presented on the left. A few overlapping points have been slightly adjusted for presentation purposes. Panels a–d illustrate examples of mires in different degradation states (Classes 1–4), the locations of which are also indicated on the map (a–d). Map data: waterbodies (Finnish Environment Institute, 2021); basemap and borders (Runfola et al., 2020); permafrost distribution (Brown et al., 2002). Photos: Oona Leppiniemi, 2025.

The landscape of the region is characterized by varying topography created by low-lying mountains and valleys. Study area belongs to the sub-Arctic vegetation zone, and more specifically, to the northern oroboreal and orohemiarctic zones (Ahti et al. 1968; Hämet-Ahti 1963). Vegetation in the region ranges from open mires and fens in the valley bottoms to mountain birch forest and treeless tundra on mountains. Sporadic and discontinuous permafrost is found in the study area above the treeline (about 400 m asl) and in peatlands, where it is protected by the peat layers (Gisnås et al., 2017; Seppälä, 2011). In general, the permafrost in Finnish palsa mires is less than 1000 years old and the latest documentations of new palsas forming in Finland date back to early 2000s (Oksanen, 2008; Seppälä, 2005). Permafrost degradation is the ongoing trend in Finland and a recent study reported that the area occupied by palsas has decreased 77 to 90 % in two mires located in our study area since 1959 (Verdonen et al., 2023).

### 2.2. Vegetation survey

In August 2025, we surveyed the plant communities of 14 mires representing different stages of permafrost degradation (Fig. 1). The mires were selected based on the classification of palsa mire degradation states by Ruuhijärvi et al. (2022). To supplement the vegetation survey of collapsed palsa mires, we included two mires from the dataset of Tammilehto et al. (2024) due to their easier accessibility. Ruuhijärvi et al. (2022) classified 282 Finnish palsa mires into five classes by interpretating their degradational state from orthoimages and by conducting validating field visits. They applied the classes defined by Kontula & Raunio (2019) which are also used in the assessment of threatened habitat types in Finland. We adopted the same classification in our study but reversed the class numbering to more intuitively describe the progressive permafrost degradation in palsa mires. In addition, we combined the two classes showing least signs of permafrost thaw. The final classification of the degradational states in this study was as follows:

- **Class 1**: Near pristine mires: < 30 % of palsas thawed, palsas are still in good condition (Fig. 1a)
- **Class 2**: Notably degraded mires: 30–60 % of palsas thawed, some of the palsas are still in good condition, thermokarsts are present (Fig. 1b)
- **Class 3**: Highly degraded mires: > 60 % of palsas thawed, thermokarsts and thaw circles are present (Fig. 1c)
- **Class 4**: Collapsed mires: palsas have collapsed, only individual palsa remnants may be present but otherwise unfrozen fen prevail (Fig. 1d).

We surveyed plant communities along transects that spanned from the tops of the palsas to the surrounding unfrozen fen. Thus, the transects represented water table depth gradient from hummock level to flark level, and alkalinity gradient from ombrotrophic bog to minerotrophic fen (Laitinen et al., 2008; Økland et al., 2001; Sjörs, 1948; Fig. 2). Each transect consisted of six 50 cm x 50 cm vegetation plots that were positioned to representative microhabitats within each palsa mires. The first two plots were placed on top of palsas, representing mostly ombrotrophic bog vegetation, more specifically *Rubo chamaemori-Dicranion elongati* alliance (Jiroušek et al., 2022). They are dry and windswept microhabitats, influenced by the permafrost core. Typically, the vegetation on these palsa tops consists of lichens, mostly dominated by different *Cladonia* species, and dwarf shrubs, such as *Empetrum nigrum*, *Andromeda polifolia*, and *Vaccinium* species. In addition, *Betula nana* and *Rubus chamaemorus* are common (Ruuhijärvi, 1960; Zuidhoff & Kolstrup, 2005; Fig. 2A). The next two study plots of the transect were positioned on the palsa slopes: one on the upper slope and one on the lower slope. Palsa slopes, experiencing contrasting snow and shading conditions, act as a small-scale ecotone between the palsa top and the adjacent fen, and therefore, the plant communities of the slopes can share species from both microhabitats (Fig. 2B). The final two plots of the transect were placed on the unfrozen fen, representing mostly flark level with some lawn communities, ranging from poor to rich fen. Fen microhabitat is characterized by *Carex*, *Eriophorum*, and *Sphagnum* species (Errington et al. 2024; Ruuhijärvi, 1960; Fig. 2C).

**Figure 2.**
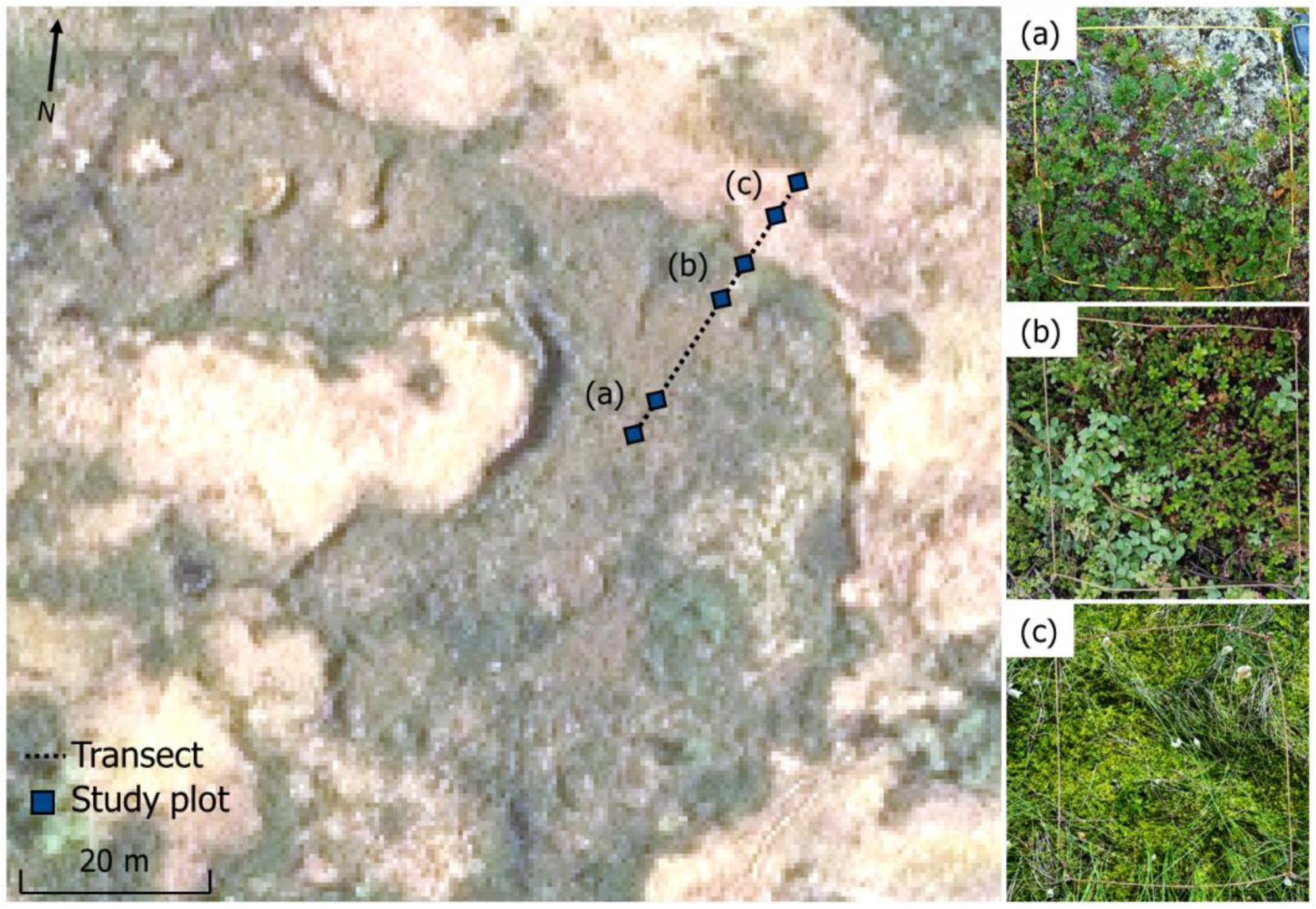
A vegetation transect across a palsa in the Karjalankoski mire (Class 2; 68.822081° N, 21.205174° E) is shown on the left to illustrate the vegetation survey design. Along the transect, six 50 x 50 cm study plots were placed on three different microhabitats: two on the palsa top (a), two on the slope (b) and two on the unfrozen fen (c). On the right are examples of the study plots illustrating the typical vegetation of each microhabitat (a–c). Orthoimage is from the National Land Survey of Finland, 2021. Photos: Oona Leppiniemi, 2025.

Plots within the same microhabitats were placed approximately 1 m apart, however, on the slopes, the spacing of the plots varied (1.0–11.0 m) to capture conditions across the upper and lower slopes. For collapsed mires (Class 4), we established the transects using the same logic by estimating the historical extent of the palsas. Our estimation was based on the historical orthoimages that we obtained from the National Land Survey of Finland, which we combined with our assessments of the mire morphology, vegetation, and moisture patterns in the field.

We recorded the cover of each vascular plant, bryophyte, and lichen species from each study plot using a percentage cover scale (< 1, 1, 2, 3, 4, 5, 7, 10, 20, … 90, 95, 96, 97, 98, 99, 100 %). In general, a total cover value was given to both liverworts (*Hepaticae*) and crustaceous lichen species, but the coverages of some common and distinctive species, such as liverwort *Ptilidium ciliare* and crustaceous lichen *Ichmadophila ericetorum*, were estimated separately. Specimens of bryophytes and lichens that we could not identify in the field were collected and subsequently identified in the lab. Some of the specimens we were able to identify only at genus level. For analyses, we classified vascular plant species into six different growth forms based on generally used classification for Arctic-alpine species (e.g., Chapin et al., 1996), and bryophytes and lichens both into five different growth forms based on Lett et al. (2022) and Nelson et al. (2015), respectively. The full list of identified taxa and different growth forms is provided in the Supplementary Materials (Tables S1– S2). In addition, we estimated the overall vegetation cover for all species combined, and for vascular plants, bryophytes, and lichens separately. The survey included a total of 33 transects (Class 1: 10 transects from four mires; Class 2: 8 transects from four mires; Class 3: 8 transects from three mires; Class 4: 7 transects from three mires) and a total of 198 study plots.

### 2.3. Statistical analysis

We studied the shifts in plant community composition across all and in different microhabitats in four distinctive stages of permafrost degradation (Classes 1–4) by using non-metric multidimensional scaling (NMDS). NMDS was calculated with absolute cover values of species (%) and based on Bray-Curtis dissimilarities. Plant growth forms were fitted as correlation vectors into the ordination. NMDS was also performed separately for each class to consider the within class variation in plant community composition of the studied mires. We used permutational manova (PERMANOVA; Anderson, 2001) with 999 permutations to test whether permafrost degradation class explained the variation in the plant community composition across all microhabitats and within each microhabitat. The permutations were constrained within study mires using a block design to account for the hierarchical structure of the data. Both analyses were performed in R version 4.5.2 (R Core Team, 2025) using ‘vegan’ package version 2.7.2 (Oksanen et al., 2024).

We used indicator species analysis to further explore the compositional differences and to identify key species assemblages co-occurring at each stage of permafrost degradation. To do this we utilized ‘indicspecies’ package version 1.8.0 in R (Cáceres & Legendre, 2009). Indicator assemblages consisting of species pairs were calculated for each permafrost degradation class and separately for each microhabitat and degradation class, by using function ‘indicators’ with alpha set at standard value of 0.05. We applied thresholds of 0.6 for specificity and 0.4 for fidelity to identify statistically significant indicator assemblages for each permafrost degradation class. For analyses focusing on different microhabitats, slightly lower thresholds (0.4 for specificity and 0.2 for fidelity) were used to allow detection of significant indicator assemblages within each class.

To quantify the impact of permafrost degradation to the plant diversity in palsa mires we explored the alpha, beta, and gamma diversity (Whittaker, 1960, 1972). Alpha diversity was quantified at microhabitat and plot levels. At microhabitat levels, we calculated the number of different species separately for each microhabitat and taxonomic group (i.e., vascular plants, bryophytes, and lichens) at each stage of permafrost degradation (Classes 1–4). At the plot-level, we calculated species richness and the effective number of common and dominant species. We calculated these measures for all species groups separately and for all groups combined. To compute the effective number of common and dominant species, we used Hill numbers q = 1 (i.e. Shannon entropy) and q = 2 (i.e. inverse Simpson) in ‘hillR’ packace version 0.5.2 (Li, 2018). These values were then analyzed in generalized linear mixed models (GLMMs), in which we used microhabitats, permafrost degradation class, and their interaction as explanatory variables. The study site was incorporated into the models as random effect to account for non-independence of our study plots. For lichens the models were fitted only for palsa tops and slopes, as they were not found in fens. GLMMs were fitted using ‘glmmTMB’ package version 1.1.14 (Brooks et al., 2017), and residuals were evaluated with ‘DHARMa’ package version 0.4.7 (Hartig, 2024). In cases where residual diagnostics indicated problems, we used logarithmic transformations to improve the model fit.

Beta diversity was quantified as a multivariate dispersion (distance to spatial median) based on Bray-Curtis dissimilarities by using ‘betadisper’ in ‘vegan’ package. Differences in multivariate dispersion among permafrost degradation classes were tested using permutation tests (999 permutations), applying the same restricted permutation scheme (block design) as in the PERMANOVA. To further explore within-class variation, we assessed beta diversity patterns across different microhabitats by calculating their plot-level distances to the spatial median.

Lastly, we estimated the relevance of vascular plant communities in different microhabitats and degradation classes to other biodiversity, as well as their contribution to a pivotal ecosystem function, nectar production. To do this, we used species-specific indicator values for biodiversity relevance and nectar production produced by Tyler et al. (2021). Biodiversity relevance is a coarse estimate of the number of associated organisms associated with each plant species. It uses a logarithmic scale ranging from 1 (< 6 associated species) to 8 (> 400 species). Nectar production is an approximation of the species-specific rate of nectar production (g sugar/m^2^/year), using a logarithmic scale with values ranging from 1 (no nectar production) to 7 (very large; > 200 g) (Tyler et al., 2021). Indicator values were found for all vascular plant species in the data. Using the cover estimates as weights, we first calculated community weighted means of both indicators for each plot, using the package ‘FD’ version 1.0–12.3 (Laliberté et al. 2025). We then used GLMMs to analyze the effect of microhabitat, degradation class and their interactions to community weighted means of both indicators, with site as a random effect. Residuals of the models were checked with ‘DHARMa’ package.

## 3. Results

### 3.1. Plant communities of different microhabitats

The plant community compositions varied significantly among the permafrost degradation classes in all microhabitats (Fig. 3). Plant communities of Classes 1–3 overlapped markedly in all microhabitats, while Class 4 had distinctive plant communities across the different microhabitats and when all microhabitats were combined. Top and slope communities shifted away from lichen and dwarf shrub dominated communities towards those characterized by *Sphagnum* mosses and graminoids as the permafrost completely disappears (Class 4) (Fig. 3ab, Tables S2–S3). Fen communities of Classes 1– 3 were characterized by sedges, whereas those of Class 4 were more associated with forbs and bryophytes with branched shoots (Fig. 3c). Among collapsed mires (Class 4), the direction of compositional shift from collapsed top to fen communities was different for each mire and indicated that the collapsed top communities did not resemble the surrounding fen community (Fig. S1; Table S4).

**Figure 3.**
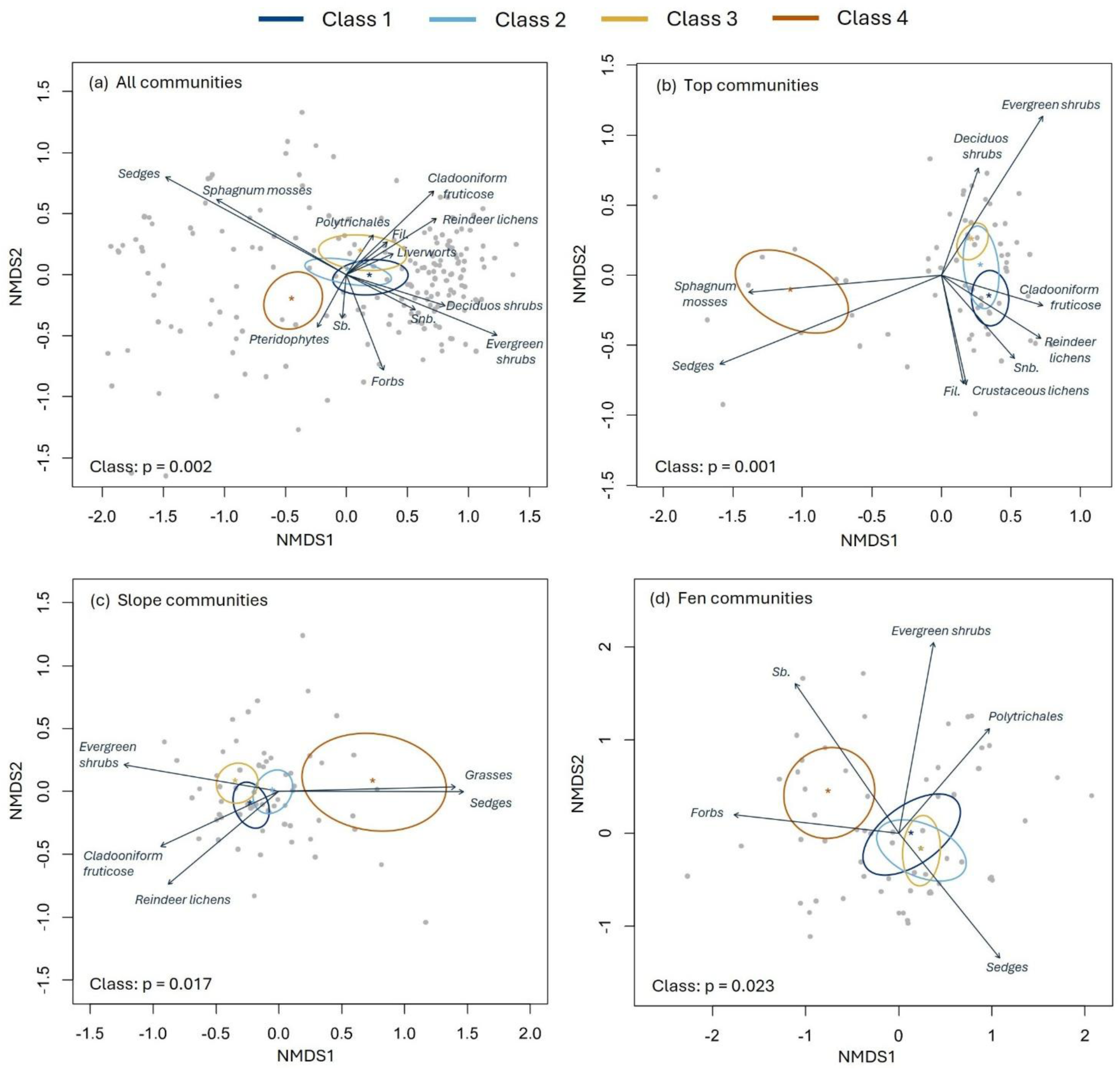
Non-metric multidimensional scaling (NMDS) illustrating the shifts in vegetation community compositions of different microhabitats of palsa mires in different stages of permafrost degradation (Classes 1–4) combined and separately (a–d). Ellipses illustrate 95 % confidence intervals of group centroids, and the significance of permafrost degradation classes is derived from PERMANOVA. Arrows represent significant (p < 0.05) correlation vectors for the cover (%) of different vascular plant, bryophyte, and lichen growth forms (see Table S3 for r^2^ and p-values of the vectors). Abbreviated growth forms refer to sprawling filamentous fruticose (Fil.), bryophytes with branched shoots (Sb.), and bryophytes with sparsely branched or unbranched shoots (Snb.).

### 3.2. Indicator species assemblages of different microhabitats

Indicator species analysis revealed distinctive indicator species assemblages for different permafrost degradation classes (Table 1). The indicator assemblages (n = 11) of Class 1 mires consisted of nine different species, and each indicator pair included *Flavocetraria nivalis*, *Andromeda polifolia*, or some of the *Cladonia* species. For Class 2, the only statistically significant indicator pair included *A. polifolia* and *Polytrichum strictum*. For Class 3 ten significant indicator pairs were recognized, and they included ten different species in total. However, *Ptilidium ciliare* was included in eight of the indicator pairs. *Betula nana* and *Eriophorum vaginatum* formed the only significant indicator pair for Class 4 (Table 1).

**Table 1.** Indicator assemblages of species pairs (p < 0.05) for the plant communities of each permafrost degradation class (Classes 1–4). ‘A’ refers to specificity, which describes the degree to which an indicator assemblage is restricted to a given degradation class, while ‘B’ refers to fidelity describing how consistently the indicator occurs within that class. ‘sqrtIV’ is the square root of the indicator value, summarizing overall indicator strength by combining A and B.

|  | Indicator assemblage | A | B | sqrtIV | p-value |
| --- | --- | --- | --- | --- | --- |
| **Class 1** | 1. *Cladonia arbuscula* + *Flavocetraria nivalis* | 0.852 | 0.467 | 0.630 | 0.005 |
|  | 2. *F. nivalis* + *Rubus chamaemorus* | 0.691 | 0.450 | 0.557 | 0.005 |
|  | 3. *Andromeda polifolia* + *C. arbuscula* | 0.637 | 0.467 | 0.545 | 0.005 |
|  | 4. *Cladonia gracilis* + *F. nivalis* | 0.763 | 0.383 | 0.541 | 0.005 |
|  | 5. *Empetrum nigrum* + *F. nivalis* | 0.644 | 0.433 | 0.528 | 0.005 |
|  | 6. *A. polifolia* + *C. gracilis* | 0.702 | 0.367 | 0.507 | 0.005 |
|  | 7. *Dicranum elongatum* + *F. nivalis* | 0.833 | 0.300 | 0.500 | 0.005 |
|  | 8. *A. polifolia* + *Cladonia pleurota* | 0.701 | 0.333 | 0.483 | 0.005 |
|  | 9. *C. pleurota* + *F. nivalis* | 0.623 | 0.333 | 0.456 | 0.020 |
|  | 10. *A. polifolia* + *D. elongatum* | 0.636 | 0.300 | 0.437 | 0.010 |
|  | 11. *A. polifolia* + *Flavocetraria cuculata* | 0.610 | 0.300 | 0.428 | 0.025 |
| **Class 2** | 1. *A. polifolia* + *Polytrichum strictum* | 0.632 | 0.333 | 0.459 | 0.010 |
| **Class 3** | 1. *Betula nana* + *Ptilidium ciliare* | 0.945 | 0.479 | 0.673 | 0.005 |
|  | 2. *P. ciliare* + *Vaccinium vitis-idaea* | 0.963 | 0.479 | 0.664 | 0.005 |
|  | 3. *E. nigrum* + *P. ciliare* | 0.860 | 0.479 | 0.642 | 0.005 |
|  | 4. *E. nigrum* + *V. vitis-idaea* | 0.635 | 0.583 | 0.609 | 0.005 |
|  | 5. *Cladonia gracilis* + *P. ciliare* | 0.839 | 0.417 | 0.609 | 0.005 |
|  | 6. *C. arbuscula* + *P. ciliare* | 0.881 | 0.375 | 0.575 | 0.005 |
|  | 7. *Cladonia rangiferina* + *P. ciliare* | 0.858 | 0.375 | 0.567 | 0.005 |
|  | 8. *Dicranum fuscenscens* + *P. ciliare* | 0.947 | 0.313 | 0.544 | 0.005 |
|  | 9. *P. ciliare* + *R. chamaemorus* | 0.776 | 0.375 | 0.540 | 0.005 |
|  | 10. *Cladonia sulphurina* + *E. nigrum* | 0.606 | 0.396 | 0.490 | 0.005 |
| **Class 4** | 1. *B. nana* + *Eriophorum vaginatum* | 0.645 | 0.310 | 0.447 | 0.005 |

Additional analyses made for different microhabitats showed that top communities of Classes 1–3 were most often indicated by different *Cladonia* and *Flavocetraria* species, but *P. strictum* was also common in Class 2 indicators, and *P. ciliare* in Class 3 (Table S5). Indicator assemblages for top communities of collapsed mires (Class 4) varied more, but included species such as *Vaccinium oxycoccos*, *Eriophorum vaginatum*, and *Sphagnum russowii*. Slope communities across all degradation classes were best indicated by different bryophytes and dwarf shrubs, when compared to lichen dominated top communities. Indicator assemblages for fen communities consisted mostly of different *Carex*, *Eriophorum*, and *Sphagnum* species (Table S5).

### 3.3. Alpha and beta diversity of degrading palsa mires

In total, 122 different taxa were identified in our study plots, the number of vascular plant, bryophyte, and lichen taxa being 39, 54, and 29, respectively (Fig. 4a; Table S1). When the species richness of different taxonomic groups was examined along the permafrost degradation classes, only lichens showed clear decline from Class 1 with 26 taxa to Class 4 with only four taxa present. On contrast, vascular plant and bryophyte richness were highest in Class 4. At the microhabitat level, palsa slopes sustained the highest species richness when all classes and taxa were considered together (Fig. 4b). In top communities, species richness was only slightly higher in Classes 1 and 2, whereas in slope communities, species richness was higher in Classes 1 and 4. Species richness in fen communities decreased steadily from Class 1 to 3 but increased markedly, and was highest, in Class 4 (Fig. 4b).

**Figure 4.**
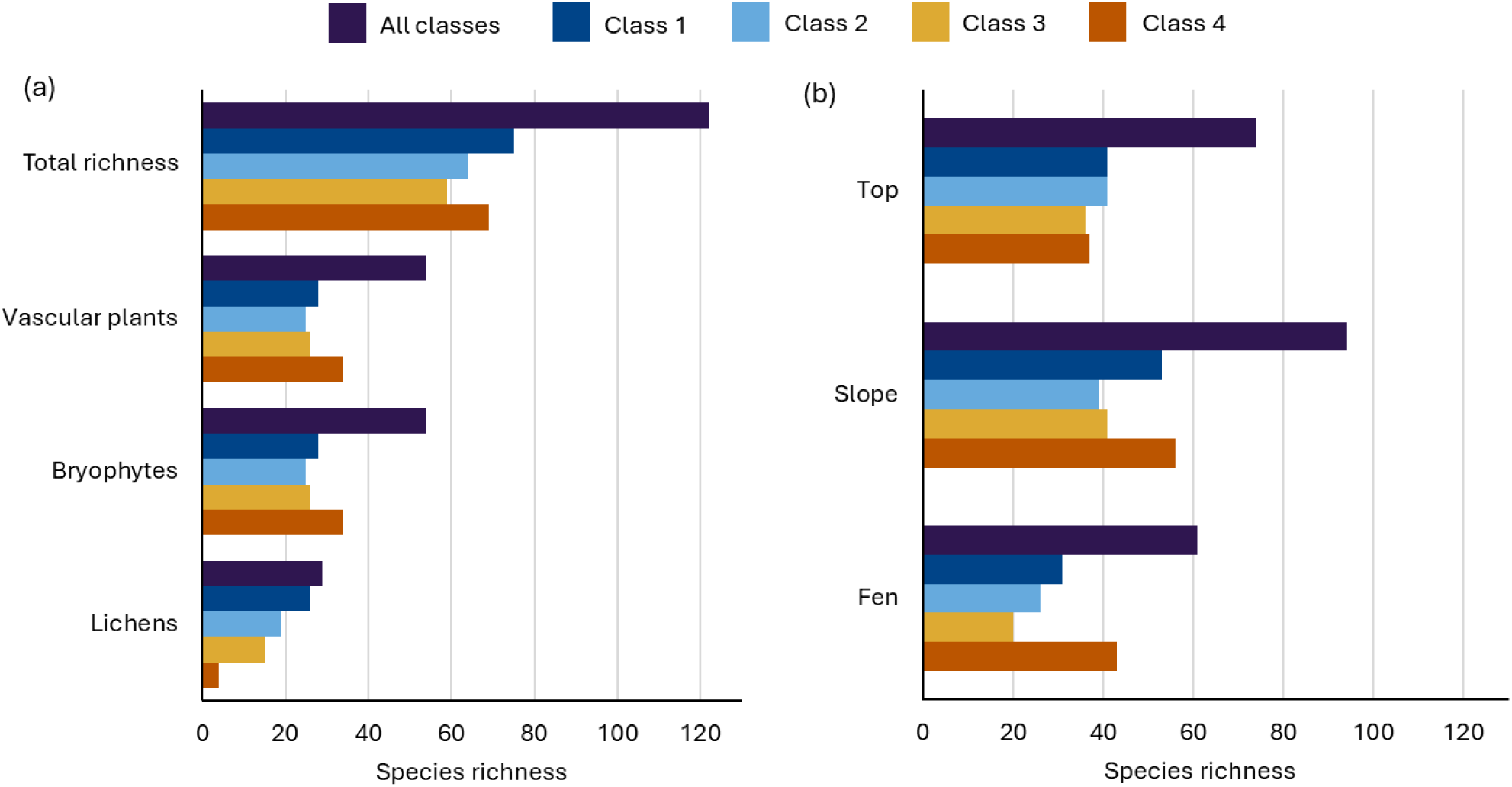
Species richness (the number of different taxa) across the permafrost degradation classes for all taxonomic groups together and separately for each group (a). Species richness across all classes and within each permafrost degradation class in the microhabitats studied (b).

The plot-level species richness analysis based on GLMMs showed that, in general, the mean species richness was greater in palsa tops and slopes than in fen habitats across all permafrost degradation classes and taxonomic groups (excluding lichens that were not present in fens) (Fig. 5). However, some variation in species richness patterns was observed among classes and the taxonomic groups. Palsa tops in Class 4 exhibited the lowest mean total richness, whereas in fens Class 4 had the highest mean richness among classes (Fig. 5a). Mean vascular plant and bryophyte richness were also highest in slopes of Class 4 (Fig. 5b–c). In contrast, mean vascular plant richness was lowest in Class 3 fens, while mean bryophyte richness in palsa tops was lowest in Class 1. The most distinct, yet expected, plot-level pattern was the near complete loss of lichens in Class 4 (Fig. 5d). Microhabitat was a highly significant variable (p < 0.001) in all models, whereas the importance of class and the microhabitat × class interaction varied among the models fitted for the different taxonomic groups (Fig. 5). GLMMs for the effective number of common and dominant species reflected similar patterns to species richness (see Figs. S2–S3).

**Figure 5.**
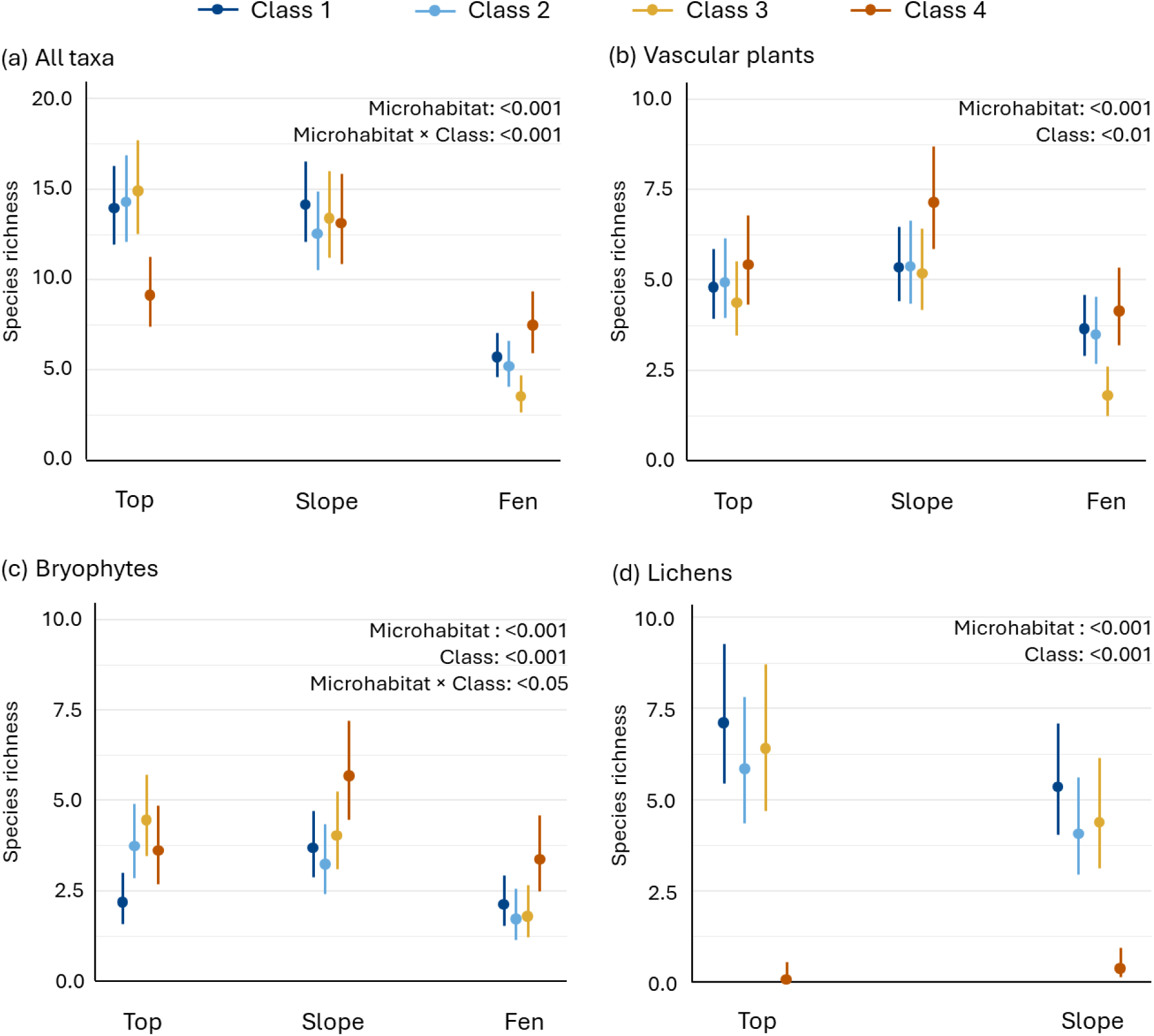
Plot-level mean species richness with 95 % confidence intervals for all taxonomic groups combined (a) and for each group separately (b–d) across the studied microhabitats. Results are based on generalized linear mixed models (GLMMs), and statistically significant explanatory variables are indicated in each panel. Note the different y-axis scale in panel (a).

Beta diversity (i.e. distance to spatial median) varied significantly among the permafrost degradation classes (p < 0.001), with Classes 1–3 showing the greatest within-class dispersion (Fig. 6a). In contrast, Class 4 displayed lower within-class dispersion but the greatest distance to spatial median, reflecting overall higher beta diversity. Microhabitat patterns of top and slope communities were consistent with this trend. However, in fen communities, Class 4 had similar within-class variation and distance to spatial median compared to Classes 1–3 (Fig. 6b).

**Figure 6.**
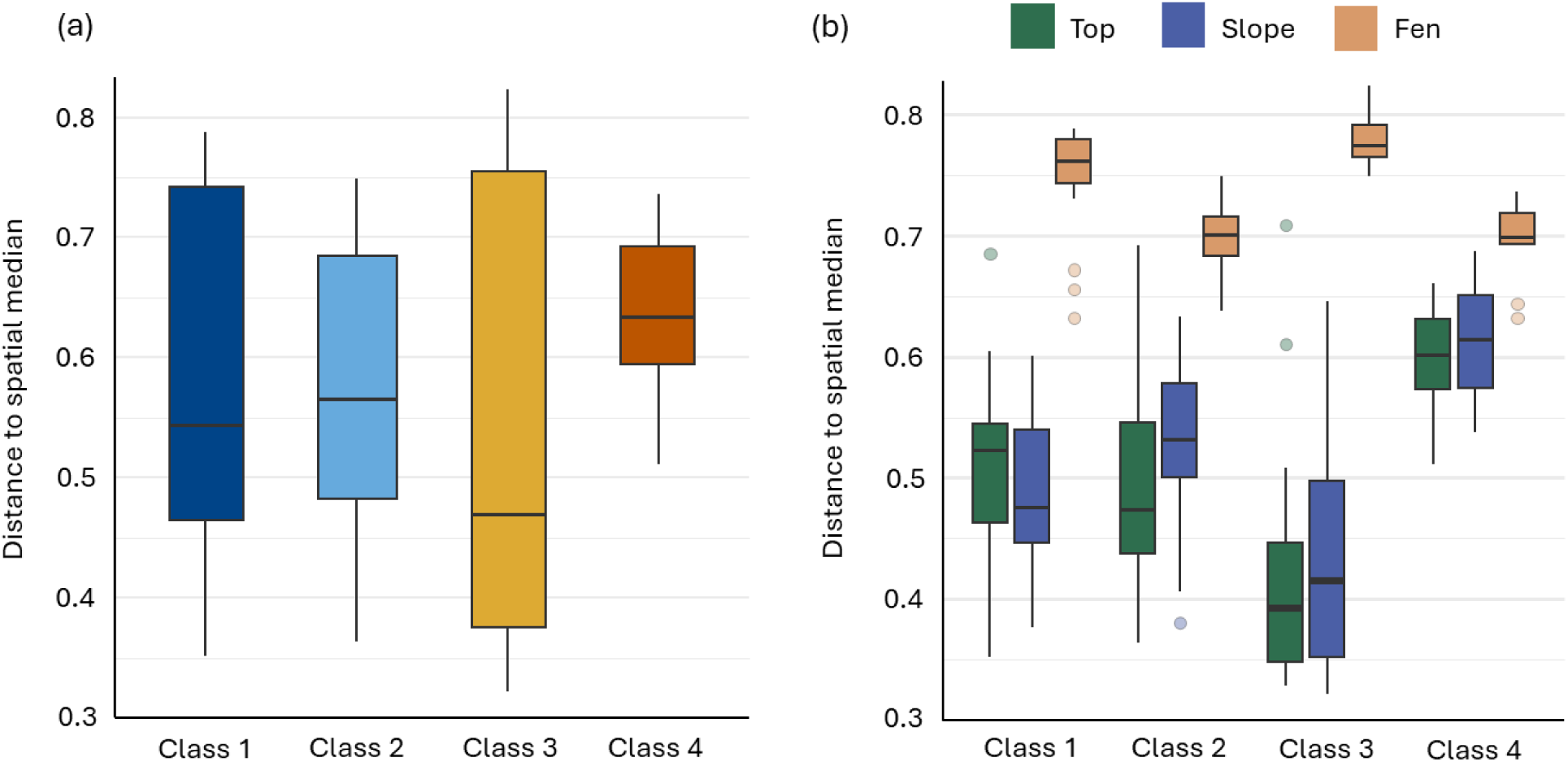
Beta diversity (distance to spatial median) across the permafrost degradation classes (a), and across the different microhabitats (b).

### 3.4. Biodiversity relevance and nectar production

In general, biodiversity relevance of vascular plants was highest on the palsa slopes (Fig. 7a), which also showed minimal variation among classes. This coincides with the highest absolute and relative coverage of vascular plants on palsa slopes across different microhabitats (Table S6). In top habitats, Class 4 had markedly lower biodiversity relevance when compared to other classes. In contrast, the biodiversity relevance was highest in fen habitats of Class 4. Nectar production reflected similar patterns to biodiversity relevance, with lowest values (i.e., nearly nonexistent production) associated with fen habitat (Fig. 7b). Classes 3 and 4 showed clearly lower nectar production values on top of palsas when compared to Classes 1 and 2. All variables used in the nectar production model were statistically significant (p < 0.001), while the permafrost degradation class was not statistically significant variable in the model for biodiversity relevance (Fig. 7).

**Figure 7.**
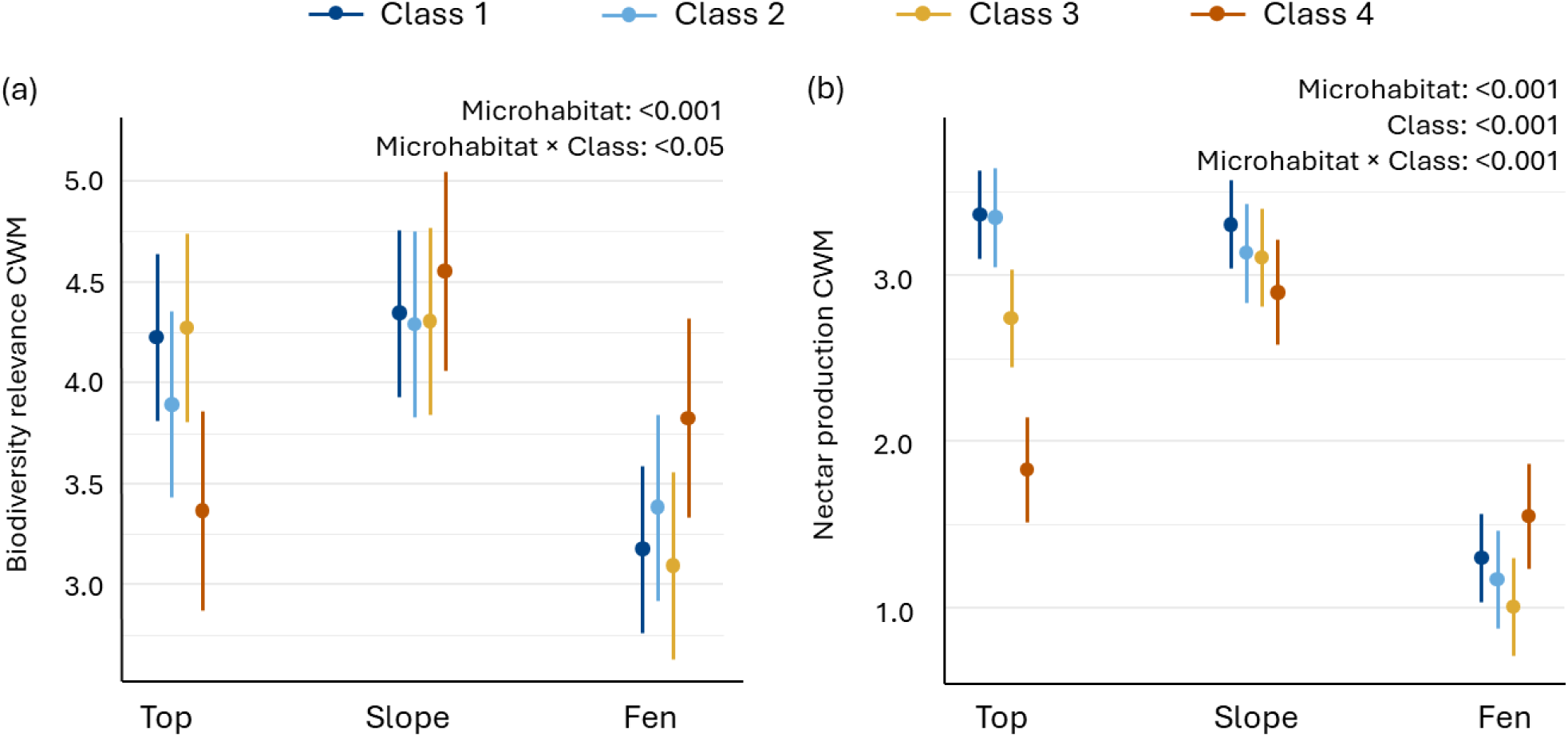
Generalized linear mixed model (GLMM) results for community weighted means (CWM) of biodiversity relevance (a) and nectar production (b) with 95 % confidence intervals. Models were run for vascular plants only. Absolute and relative covers of vascular plants are provided in Supplementary Materials in Table S6. Values for Biodiversity relevance: 3 = 13–24 associated species, 4 = 25–50, 5 = 51–100. Values for Nectar production: 1 = No nectar production (0 g sugar/m^2^/year), 2 = Nectar production insignificant (< 0.2 g), 3 = Nectar production small (0.2–5 g), 4 = Nectar production modest (5–20 g) (Tyler et al., 2021).

## 4. Discussion

Here we quantified the impact of permafrost degradation on plant community composition and diversity of vascular plants, bryophytes, and lichens by surveying vegetation of 14 palsa mires located in northern Finland. We show that any presence of permafrost supports distinctive plant communities in palsa mires, and that the disappearance of permafrost and associated collapse of palsas lead significant shifts in the plant community composition and diversity. As we hypothesized, lichens declined strongly along the permafrost degradation gradient. However, diversity patterns of different taxonomic groups varied markedly across the microhabitats of palsa mires and different stages of permafrost degradation, and depending on the biodiversity measure. This highlights the need for inclusion of also other taxonomic groups than vascular plants and a more comprehensive assessment of biodiversity (Hillebrand et al., 2018) to understand the overall impact of permafrost degradation on the diversity of Arctic peatlands. Overall, plant community composition and the observed shifts in species richness align with previous studies from North America and Sweden (Cox et al., 2025; Errington et al., 2024; Malmer et al., 2005; Tsuyuzaki et al., 2008; Zuidhoff & Kolstrup, 2005). However, our results suggest more abrupt changes in plant community composition and a tipping point in local biodiversity as palsas collapse rather than gradual transition along permafrost degradation gradient with most rapid succession of vegetation associated with early stages of permafrost degradation (Cox et al., 2025; Errington et al., 2024).

Palsa mires underlain by any amount of permafrost (Classes 1–3) were identified to sustain distinct plant communities when compared to collapsed palsa mires where permafrost is no longer present. Our NMDS results revealed that particularly lichens and shrubs were more associated with permafrost mires, while *Sphagnum* mosses, graminoids, and sedges increased as permafrost disappears. These shifts in plant communities that we found are consistent with previous observations and modelled vegetation shifts from dry hummock vegetation towards moister vegetation and are suggested to increase in CO_2_ and CH_4_ emissions from palsa mires in future (Bosiö et al., 2012; Cox et al., 2025; Errington et al., 2024; Zuidhoff & Kolstrup, 2005).

Although the community compositions of Classes 1–3 were more resembled each other when compared to Class 4, the indicator species analysis revealed that all four permafrost degradation classes were indicated by different species assemblages. This suggests that regardless of relatively similar community composition, Classes 1–3 represent different stages of permafrost degradation. Thus, the utilized classification is robust to study the transition in plant communities and diversity in response to permafrost degradation.

More specifically, we found that the most pristine palsa mires were indicated by diagnostic species of *Rubo chamaemori-Dicranion elongati* alliance (Jiroušek et al., 2022), which are also common in dry oligotrophic tundra and mountain heaths (Haapasaari, 1988; Maliniemi et al. 2025). In Class 2 indicator assemblage, *Polytrichum strictum*, a competitive perennial species (During, 1992; van Zuijlen et al., 2023) typical for hummocks (Eurola et al., 1984, 2015) is also known for its ability to colonize disturbed, bare substrates (Groeneveld et al., 2007). The transition continues to Class 3 which we found to be often indicated by large liverwort *Ptilidium ciliare,* which is an indicator of lawn level poor fens (Eurola et al. 1984; 2015) along with typical hummock level species: low shrub *Betula nana,* and dwarf shrubs *Empetrum nigrum*, and *Vaccinium vitis-idaea*. When compared to previous classes, these species indicate somewhat moister growing conditions which are less affected by permafrost. In Class 4 indicator assemblage, the inclusion of *Eriophorum vaginatum,* a species primarily growing in low hummocks and lawn level poor fens (Eurola et al., 1984, 2015), indicates continuing change towards unfrozen fen habitats, still retaining some of the hummock characteristics of palsas.

In addition to the changes in the soil moisture conditions, the shifts in vegetation might be linked to the ability of plant roots to reach the minerogenous mire water, changes in the snow cover dynamics (Luoto et al., 2004), or to the release of nutrients and metals (Loiko et al., 2017; Patzner et al., 2022). For example, while Class 1–3 fen microhabitats were characterized by *Eriophorum angustifolium,* a common flark species that tolerates seasonally dry growing conditions (Eurola et al., 1984, 2015; Laitinen et al., 2008; Ruuhijärvi, 1960), Class 4 fens were characterized by forbs such as *Epilobium palustre* and *Comarum palustre* (Table S5), suggesting greater surface water influence and increased nutrient supply (Eurola et al., 1984; Rehell et al., 2019; Ruuhijärvi 1960). However, the potential influence of unmeasured site-specific factors cannot be ruled out.

Site-specific differences between studied mires are indicated by the high beta diversity of collapsed mires and fen microhabitats across all permafrost degradation classes. For example, most often only one to two *Sphagnum* and sedge species dominated the study plots but were different species depending on the mire, which increased beta diversity. Site-specific differences are also highlighted by high number of different species in Class 4 mires that were observed in less than five study plots (Table S2). The high beta diversity between collapsed mires may also be contributed by different timing in permafrost disappearance as the studied mires may not have reached the common endpoint in vegetation succession (Swindles et al., 2015). Historical orthoimages indicate that one of the studied mires was already permafrost-free by 2000, whereas the other two did not appear fully permafrost-free until 2012–2013. Palaeoecological studies, vegetation resurveys and particularly long-term monitoring of the plant communities of palsa mires are still needed to understand the final outcomes of permafrost disappearance (Camill & Clark, 2000; Dornelas et al., 2023; Kapfer et al., 2017; Sim et al., 2021; Swindles et al., 2015).

Our results highlight the importance of including different taxonomic groups and aspects of biodiversity to understand overall change. Both showed decoupled responses along the permafrost degradation gradient, depending on the growth form or biodiversity metric, respectively. The most pristine mires sustained the highest diversity when the total species richness was calculated at class level, however collapsed mires had the highest species richness for vascular plats and bryophytes. The loss of lichens does not only reduce the local plant diversity maintained by palsa mires but has also negative impact on the availability of reindeer winter grazing areas (Ward Jones et al., 2024). Palsa slopes were recognized to sustain the highest species richness, indicating their role as micro- ecotones between palsa tops and adjacent fens. Interestingly this trend persisted across the degradation gradient, being most evident in both pristine and collapsed palsa mires. The palsa slopes of the collapsed mires were often recognizable as elevated thaw circles in the field although the former palsa tops had collapsed to the flark-level. This pattern is consistent with broader mire diversity gradients, where maximum species richness is often observed at the intersection of a relatively low water table and high nutrient availability. Similar diversity maxima have been reported during mire development (Laine et al., 2018) and along the fertility gradient of Central European fens (Hájek et al., 2006).

We assessed the potential impacts of vegetation change on other taxa indirectly, by calculating the biodiversity relevance and nectar production of vascular plant communities. The decrease in biodiversity relevance and nectar production at palsa tops and slopes of Class 4 mires is most probably linked to the reduction in coverage of different *Vaccinium* species and *Rubus chamaemorus*. These species have higher biodiversity relevance and nectar production values when compared to graminoids (Tyler et al., 2021), which are dominant vascular plants of collapsed palsa tops and fens. The shift from nectar producing species with higher biodiversity relevance to graminoids may cause ecological consequences for other taxonomic groups beyond plants, such as pollinators, and impact also local people and their livelihoods due to reduced berry yields (Ward Jones et al., 2024). However, even though biodiversity relevance and nectar production provide useful proxies, they offer only indirect evidence of broader biodiversity responses (Tyler et al., 2021). A more comprehensive understanding of the effects of permafrost degradation on palsa mire biodiversity therefore requires direct assessments of other taxonomic groups, which have so far remained scarce (however, see Järvinen & Sammalisto, 1976; Markkula, 2014).

Overall, our study demonstrated that permafrost degradation has significant impacts on the biodiversity of palsa mires. Although our data does not imply a risk for specific threatened plant species, instead, a unique peatland vegetation type, with a clear positive impact on local biodiversity, will be lost if the permafrost degradation continues and palsa mires become permafrost free. Considering the current rate of climate warming in the Arctic (Rantanen et al., 2022), the widespread degradation of palsa mires across the Northern Hemisphere (Borge et al., 2017; Mamet et al., 2017; Olvmo et al., 2020; Wang et al., 2023), and predictions indicating loss of suitable environmental spaces for palsa mires by the 21^st^ century (Aalto et al., 2014, 2017; Fewster et al., 2022; Leppiniemi et al., 2023) this threat to local biodiversity can be considered both substantial and likely accelerating under ongoing climate change. Given the observed biodiversity tipping point associated with the palsa collapse, there is clearly an urgent need for expanded research efforts on these critically endangered habitats across the Northern Hemisphere. Vegetation surveys, including multiple different taxonomic groups and regions, will provide essential baseline data for biodiversity monitoring and conservation.

## Acknowledgements

Oona Leppiniemi acknowledges the funding from the Emil Aaltonen Foundation, Tuija Maliniemi from Research Council of Finland (project 370154), Julia Kemppinen thanks the Research Council of Finland funding (grant no. 353218; 370245), and Mika Jokikokko the Kone Foundation (grant no. 202303348). We also want to acknowledge the Kilpisjärvi Biological Station for providing accommodation and research facilities throughout the fieldwork.

## Author contributions

Oona Leppiniemi and Tuija Maliniemi conceived the ideas and designed the methodology. Julia Kemppinen supported the conceptualization of the study, while Mika Jokikokko contributed to the methodology. Oona Leppiniemi collected the data with support from Tuija Maliniemi. Oona Leppiniemi and Tuija Maliniemi analyzed the data. Oona Leppiniemi led the writing of the manuscript and did data visualizations, with contributions from Tuija Maliniemi. Mika Jokikokko and Julia Kemppinen reviewed and edited drafts proving important inputs. All authors contributed critically to the manuscript and gave final approval for publication.

## Data Availability Statement

The research data and code supporting this study have been deposited in Zenodo (https://doi.org/10.5281/zenodo.22227494). During peer review, access to the repository is restricted and provided to reviewers via a private link. The data and code will be made publicly available upon acceptance for publication.

