## Supplementary Material for "Intertwined degradation of plant diversity and permafrost in sub-Arctic palsa mires"

**Table S1.** List of the vascular plant, bryophyte, and lichen taxa that were identified. The abbreviation of the plant growth form of the taxa is in parenthesis. Vascular plants were grouped according to the classification used in the tundra (see e.g., Chapin et al. (1996), bryophytes as in Lett et al. (2022), and lichens as in Nelson et al. (2015).*

| **Vascular plants** (n = 39) | **Bryophytes** (n = 54) | **Lichens** (n = 29) |
|---|---|---|
| *Andromeda polifolia* (EVE) | *Aulacomnium palustre* (SNB) | *Cetraria ericetorum* (FIL) |
| *Arctous alpina* (DEC) | *Barbilophozia lycopodioides* (LIV) | *Cetraria islandica* (FIL) |
| *Betula nana* (DEC) | *Barbilophozia* spp. (LIV) | *Cetraria muricata* (FIL) |
| *Betula pubescens* ssp. *Czerepanovii* (DEC) | *Calliergon cordifolium* (SB) | *Cladonia amaurocrae* (CLA) |
| *Calamagrostis lapponica* (GRA) | *Cinclidium subrotundum* (SNB) | *Cladonia arbuscula* (REIN) |
| *Calamagrostis neglecta* (GRA) | *Cynodontium* spp. (SNB) | *Cladonia carneola* (CLA) |
| *Calamagrostis phragmitoides* (GRA) | *Dicranella cerviculata* (SNB) | *Cladonia chlorophaea* (CLA) |
| *Carex acuta* (SED) | *Dicranella crispa* (SNB) | *Cladonia coccifera* (CLA) |
| *Carex aquatilis* (SED) | *Dicranella* spp. (SNB) | *Cladonia cornuta* (CLA) |
| *Carex canescens* (SED) | *Dicranum drummondii* (SNB) | *Cladonia digita* (CLA) |
| *Carex chordorrhiza* (SED) | *Dicranum elongatum* (SNB) | *Cladonia fimbriata* (CLA) |
| *Carex limosa* (SED) | *Dicranum fuscenscens* (SNB) | *Cladonia furcata* (CLA) |
| *Carex paupercula* (SED) | *Dicranum scoparium* (SNB) | *Cladonia gracilis* (CLA) |
| *Carex rostrata* (SED) | *Dicranum undulatum* (SNB) | *Cladonia macrophylla* (CLA) |
| *Carex rotundata* (SED) | *Ditrichum* spp. (SNB) | *Cladonia pleurota* (CLA) |
| *Cerastium fontanum* (FORB) | *Hepaticae* spp. (LIV) | *Cladonia rangiferina* (REIN) |
| *Comarum palustre* (FORB) | *Mylia anomala* (LIV) | *Cladonia squamosa* (CLA) |
| *Empetrum nigrum* (EVE) | *Oligotrichum hercynicum* (SNB) | *Cladonia stellaris* (REIN) |
| *Epilobium palustre* (FORB) | *Oncophorus* spp. (SNB) | *Cladonia subulata* (CLA) |
| *Equisetum fluviatile* (PTER) | *Oncophorus wahlenbergii* (SNB) | *Cladonia sulphurina* (CLA) |
| *Equisetum palustre* (PTER) | *Paludella squarrosa* (SNB) | *Cladonia uncialis* (CLA) |
| *Eriophorum angustifolium* (SED) | *Pleurozium schreberi* (SB) | *Crustaceous* lichens (CRU) |
| *Eriophorum vaginatum* (SED) | *Pohlia nutans* (SNB) | *Flavocetraria cuculata* (FIL) |
| *Galium palustre* (FORB) | *Polytrichum hyperboreum* (POL) | *Flavocetraria nivalis* (FIL) |
| *Huperzia selago* (PTER) | *Polytrichum juniperinum* (POL) | *Gowardia nigricans* (FIL) |
| *Menyanthens trifoliata* (FORB) | *Polytrichum strictum* (POL) | *Ichmadophila ericetorum* (CRU) |
| *Poa* sp. (GRA) | *Pseudobryum cinclidioides* (SNB) | *Ochrolechia frigida* (CRU) |
| *Rhododendron tomentosum* (EVE) | *Ptilidium ciliare* (LIV) | *Ochrolechia* spp. (CRU) |
| *Rubus chamaemorus* (FORB) | *Ptychostomum pseudotriquetrum* (SNB) | *Peltigera aphthosa* (FOL)** |
| *Salix lapponum* (DEC) | *Ptychostomum* spp. (SNB) | |
| *Salix myrtilloides* (DEC) | *Rhizomnium punctatum* (SNB) | |
| *Salix phylicifolia* (DEC) | *Sarmentypnum exannulatum* (SB) | |
| *Spinulum annotinum* (PTER) | *Sarmentypnum sarmentosum* (SB) | |
| *Trichophorum cespitosum* (SED) | *Sarmentypnum tundrae* (SB) | |
| *Vaccinium microcarpum* (EVE) | *Scorpidium cossonii* (SB) | |
| *Vaccinium myrtillus* (DEC) | *Scorpidium revolvens* (SB) | |
| *Vaccinium oxycoccos* (EVE) | *Sphagnum angustifolium* (SPH) | |
| *Vaccinium uliginosum* (DEC) | *Sphagnum capillifolium* (SPH) | |
| *Vaccinium vitis-idaea* (EVE) | *Sphagnum compactum* (SPH) | |
| | *Sphagnum fallax* (SPH) | |
| | *Sphagnum fuscum* (SPH) | |
| | *Sphagnum jensenii* (SPH) | |
| | *Sphagnum lindbergii* (SPH) | |
| | *Sphagnum medium* (SPH) | |
| | *Sphagnum obtusum* (SPH) | |
| | *Sphagnum riparium* (SPH) | |
| | *Sphagnum russowii* (SPH) | |
| | *Sphagnum squarrosum* (SPH) | |
| | *Sphagnum subfulvum* (SPH) | |
| | *Sphagnum teres* (SPH) | |
| | *Stramiergon stramineum* (SB) | |
| | *Tetraplodon mnioides* (SNB) | |
| | *Warnstorfia fluitans* (SB) | |
| | *Warnstorfia pseudostraminea* (SB) | |
| | | **Total n = 122** |

*Abbreviations in Table S1 stand for DEC=deciduous shrubs, EVE = evergreen shrubs, FORB = forbs, GRA = grasses, PTER = pteridophytes, SED = sedges (vascular plants); LIV = liverworts, POL = Polytrichales, SB = shoot branched, SNB = shoots not or sparsely branched, SPH = Sphagnum (bryophytes); CLA = Cladooniform fruticose, CRU = crustaceous, FIL = sprawling filamentous fruticose, FOL = foliose, and REIN = reindeer lichens (lichens). **FOL was excluded from the analyses as it was recorded only once.

**Table S2.** List of the vascular plant, bryophyte, and lichen taxa that were identified in each permafrost degradation class. The percentage of plots within the class (%) in which each taxon was observed is shown in parentheses, and the total number of different taxa in each class is presented in the final row.

| **Class 1** | **Class 2** | **Class 3** | **Class 4** |
|---|---|---|---|
| *Rubus chamaemorus* (71.7) | *Empetrum nigrum* (66.7) | *Betula nana* (68.8) | *Andromeda polifolia* (52.4) |
| *Empetrum nigrum* (66.7) | *Rubus chamaemorus* (66.7) | *Empetrum nigrum* (66.7) | *Eriophorum vaginatum* (50.0) |
| *Andromeda polifolia* (58.3) | *Betula nana* (58.3) | *Vaccinium vitis-idaea* (58.3) | *Betula nana* (45.2) |
| *Betula nana* (58.3) | *Vaccinium vitis-idaea* (54.2) | *Cladonia arbuscula* (54.2) | *Rubus chamaemorus* (42.9) |
| *Cladonia arbuscula* (55.0) | *Andromeda polifolia* (50.0) | *Cladonia gracilis* (54.2) | *Empetrum nigrum* (38.1) |
| *Flavocetraria nivalis* (46.7) | *Cladonia arbuscula* (50.0) | *Ptilidium ciliare* (50.0) | *Eriophorum angustifolium* (38.1) |
| *Cladonia gracilis* (45.0) | *Cladonia gracilis* (43.8) | *Rubus chamaemorus* (50.0) | *Hepaticae* spp. (38.1) |
| *Cladonia pleurota* (43.3) | *Polytrichum strictum* (37.5) | *Cladonia rangiferina* (41.7) | *Vaccinium oxycoccos* (28.1) |
| *Vaccinium uliginosum* (43.3) | *Cladonia uncialis* (35.4) | *Polytrichum strictum* (41.7) | *Polytrichum strictum* (31.0) |
| *Polytrichum strictum* (38.3) | *Cladonia pleurota* (33.3) | *Cladonia pleurota* (39.6) | *Vaccinium uliginosum* (28.6) |
| *Vaccinium vitis-idaea* (36.7) | *Vaccinium uliginosum* (33.3) | *Cladonia sulphurina* (39.6) | *Epilobium palustre* (26.2) |
| *Cladonia sulphurina* (35.0) | *Hepaticae* spp. (31.3) | *Hepaticae* spp. (39.6) | *Sphagnum riparium* (26.2) |
| *Dicranum elongatum* (33.3) | *Dicranum fuscenscens* (29.2) | *Dicranum fuscenscens* (35.4) | *Sphagnum russowii* (26.2) |
| *Cladonia rangiferina* (31.7) | *Eriophorum angustifolium* (29.2) | *Eriophorum vaginatum* (33.3) | *Carex rotundata* (23.8) |
| *Flavocetraria cuculata* (31.7) | *Flavocetraria nivalis* (29.2) | *Vaccinium uliginosum* (27.1) | *Vaccinium vitis-idaea* (21.4) |
| *Hepaticae* spp. (31.7) | *Sphagnum riparium* (27.1) | *Cladonia uncialis* (25.0) | *Sphagnum fallax* (19.0) |
| *Cladonia uncialis* (25.0) | *Cladonia rangiferina* (25.0) | *Flavocetraria cuculata* (25.0) | *Stramiergon stramineum* (19.0) |
| *Eriophorum angustifolium* (23.3) | *Flavocetraria cuculata* (25.0) | *Pohlia nutans* (22.9) | *Carex canescens* (14.3) |
| *Eriophorum vaginatum* (20.0) | *Cladonia sulphurina* (22.9) | *Sphagnum riparium* (20.8) | *Comarum palustre* (14.3) |
| *Vaccinium oxycoccos* (20.0) | *Eriophorum vaginatum* (20.8) | *Eriophorum angustifolium* (18.8) | *Pleurezium schreberi* (14.3) |
| *Dicranum fuscenscens* (18.3) | *Dicranum elongatum* (18.8) | *Flavocetraria nivalis* (16.7) | *Aulacomnium palustre* (11.9) |
| *Carex rostrata* (15.0) | *Vaccinium oxycoccos* (18.8) | *Andromeda polifolia* (12.5) | *Calliergon cordifolium* (11.9) |
| *Cetraria ericetorum* (13.3) | *Epilobium palustre* (14.6) | *Cladonia coccifera* (12.5) | *Carex aquatilis* (11.9) |
| *Cladonia coccifera* (11.7) | *Pohlia nutans* (14.6) | *Sphagnum lindbergii* (12.5) | *Carex limosa* (11.9) |
| *Polytrichum juniperium* (10.0) | *Ptilidium ciliare* (14.6) | *Pleurozium schreberi* (10.4) | *Carex rostrata* (11.9) |
| *Ptilidium ciliare* (10.0) | *Ichmadophila ericetorum* (12.5) | *Dicranum elongatum* (8.3) | *Dicranum scoparium* (11.9) |
| *Sphagnum riparium* (10.0) | *Polytrichum juniperium* (12.5) | *Dicranum drummondii* (8.3) | *Psedobryum cinclidioides* (11.9) |
| *Sphagnum russowii* (10.0) | *Carex cannescens* (10.4) | *Polytrichum juniperium* (8.3) | *Salix lapponum* (11.9) |
| *Sphagnum capillifolium* (10.0) | *Pleurozium schreberi* (10.4) | *Carex rostrata* (6.3) | *Sphagnum lindbergii* (11.9) |
| *Cladonia amaurocrae* (8.3) | *Carex aquatilis* (8.3) | *Carex rotundata* (6.3) | *Sphagnum medium* (11.9) |
| *Comarum palustre* (8.3) | *Cetraria islandica* (6.3) | *Cladonia carneola* (6.3) | *Equisetum palustre* (9.5) |
| *Vaccinium myrtillus* (8.3) | *Cladonia coccifera* (6.3) | *Cladonia squamosa* (6.3) | *Sphagnum fuscum* (9.5) |
| *Barbilophozia* spp. (6.7) | *Dicranum scoparium* (6.3) | *Cladonia stellaris* (6.3) | *Sphagnum squarrosum* (9.5) |
| *Carex aquatilis* (6.7) | *Dicranum undulatum* (6.3) | *Sphagnum angustifolium* (6.3) | *Trichophorum cespitosum* (9.5) |
| *Carex canescens* (6.7) | *Sphagnum capillifolium* (6.3) | *Vaccinium myrtillus* (6.3) | *Vaccinium myrtillus* (9.5) |
| *Pohlia nutans* (6.7) | *Sphagnum fuscum* (6.3) | *Vaccinium oxycoccos* (6.3) | *Dicranum elongatum* (7.1) |
| *Calliergon cordifolium* (6.7) | *Sphagnum squarrosum* (6.3) | *Barbilophozia lycopodioides* (4.2) | *Menyanthens trifoliata* (7.1) |
| *Cladonia fimbriata* (5.0) | *Vaccinium myrtillus* (6.3) | *Calamagrostis phragmitoides* (4.2) | *Paludella squarrosa* (7.1) |
| *Cladonia furcata* (5.0) | *Carex rostrata* (4.2) | *Cynodontium* spp. (4.2) | *Pohlia nutans* (7.1) |
| *Cladonia squamosa* (5.0) | *Carex rotundata* (4.2) | *Epilobium palustre* (4.2) | *Polytrichum juniperium* (7.1) |
| *Cladonia stellaris* (5.0) | *Cerastium fontanum* (4.2) | *Spinulum annotinum* (4.2) | *Ptilidium ciliare* (7.1) |
| *Dicranum undulatum* (5.0) | *Cladonia carneola* (4.2) | *Aulacomnium palustre* (2.1) | *Sphagnum angustifolium* (7.1) |
| *Sphagnum fallax* (5.0) | *Cladonia cornuta* (4.2) | *Betula pubescens* ssp. *Czerepanovii* (2.1) | *Sphagnum capillifolium* (7.1) |
| *Cinclidium subrotundum* (3.3) | *Cladonia subulata* (4.2) | *Carex canescens* (2.1) | *Sphagnum jensenii* (7.1) |
| *Cladonia cornuta* (3.3) | *Comarum palustre* (4.2) | *Cetraria ericetorum* (2.1) | *Sphagnum teres* (7.1) |
| *Cladonia digita* (3.3) | *Ochrolechia* spp. (4.2) | *Cladonia cornuta* (2.1) | *Vaccinium microcarpum* (7.1) |
| *Cladonia chlorophaea* (3.3) | *Sphagnum angustifolium* (4.2) | *Dicranum scoparium* (2.1) | *Warnstorfia pseudostraminea* (7.1) |
| *Dicranum scoparium* (3.3) | *Calliergon cordifolium* (2.1) | *Dicranum undulatum* (2.1) | *Calamagrostis neglecta* (4.8) |
| *Dicranella crispa* (3.3) | *Calamagrostis neglacta* (2.1) | *Dicranella* sp. (2.1) | *Carex chordorrhiza* (4.8) |
| *Epilobium palustre* (3.3) | *Carex paupercula* (2.1) | *Ditrichum* sp. (2.1) | *Carex paupercula* (4.8) |
| *Equisetum fluviatile* (3.3) | *Cladonia fimbriata* (2.1) | *Huperzia selago* (2.1) | *Cladonia arbuscula* (4.8) |
| *Ichmadophila ericetorum* (3.3) | *Cladonia squamosa* (2.1) | *Oncophorus wahlenbergii* (2.1) | *Cladonia rangiferina* (4.8) |
| *Stramiergon stramineum* (3.3) | *Cladonia stellaris* (2.1) | *Peltigera aphthosa* (2.1) | *Dicranum undulatum* (4.8) |
| *Trichophorum cespitosum* (3.3) | *Crustaceous* lichens (2.1) | *Sphagnum capillifolium* (2.1) | *Rhizomnium punctatum* (4.8) |
| *Arctous alpina* (1.7) | *Oncophorus* sp. (2.1) | *Sphagnum compactum* (2.1) | *Rhododendron tomentosum* (4.8) |
| *Aulacomnium palustre* (1.7) | *Polytrichum hyperboreum* (2.1) | *Sphagnum fuscum* (2.1) | *Salix phylicifolia* (4.8) |
| *Barbilophozia lycopodioides* (1.7) | *Ptychostonum* sp. (2.1) | *Sphagnum russowii* (2.1) | *Sarmentyonum sarmentosum* (4.8) |
| *Calamagrostis lapponica* (1.7) | *Sarmentypnum exannulatum* (2.1) | *Sphagnum subfulvum* (2.1) | *Sphagnum obtusum* (4.8) |
| *Carex rotundata* (1.7) | *Sarmentypnum tundrae* (2.1) | *Warnstorfia fluitans* (2.1) | *Carex acuta* (2.4) |
| *Cetraria islandica* (1.7) | *Sphagnum fallax* (2.1) | | *Cladonia gracilis* (2.4) |
| *Cetraria muricata* (1.7) | *Sphagnum subfulvum* (2.1) | | *Dicranum fuscenscens* (2.4) |
| *Cladonia carneola* (1.7) | *Sphagnum teres* (2.1) | | *Flavocetraria cuculata* (2.4) |
| *Cladonia macrophylla* (1.7) | *Tetraplodon mnioides* (2.1) | | *Galium palustre* (2.4) |
| *Dicranella cerviculata* (1.7) | *Vaccinium microcarpum* (2.1) | | *Mylia anomala* (2.4) |
| *Dicranella* sp. (1.7) | | | *Poa* sp. (2.4) |
| *Galium palustre* (1.7) | | | *Salix myrtilloides* (2.4) |
| *Gowardia nigricans* (1.7) | | | *Sarmentypnum exannulatum* (2.4) |
| *Ochrolechia* sp. (1.7) | | | *Scorpidium revolvens* (2.4) |
| *Ochrolechia frigida* (1.7) | | | *Scorpidium scorpioides* (2.4) |
| *Oligotrichum hercynicum* (1.7) | | | |
| *Pleurezium schreberi* (1.7) | | | |
| *Pseudobryum cinclidioides* (1.7) | | | |
| *Ptychostonum pseudodotriquetrum* (1.7) | | | |
| *Ptychostonum* sp. (1.7) | | | |
| *Sphagnum fuscum* (1.7) | | | |
| **n = 75** | **n = 64** | **n = 59** | **n = 69** |

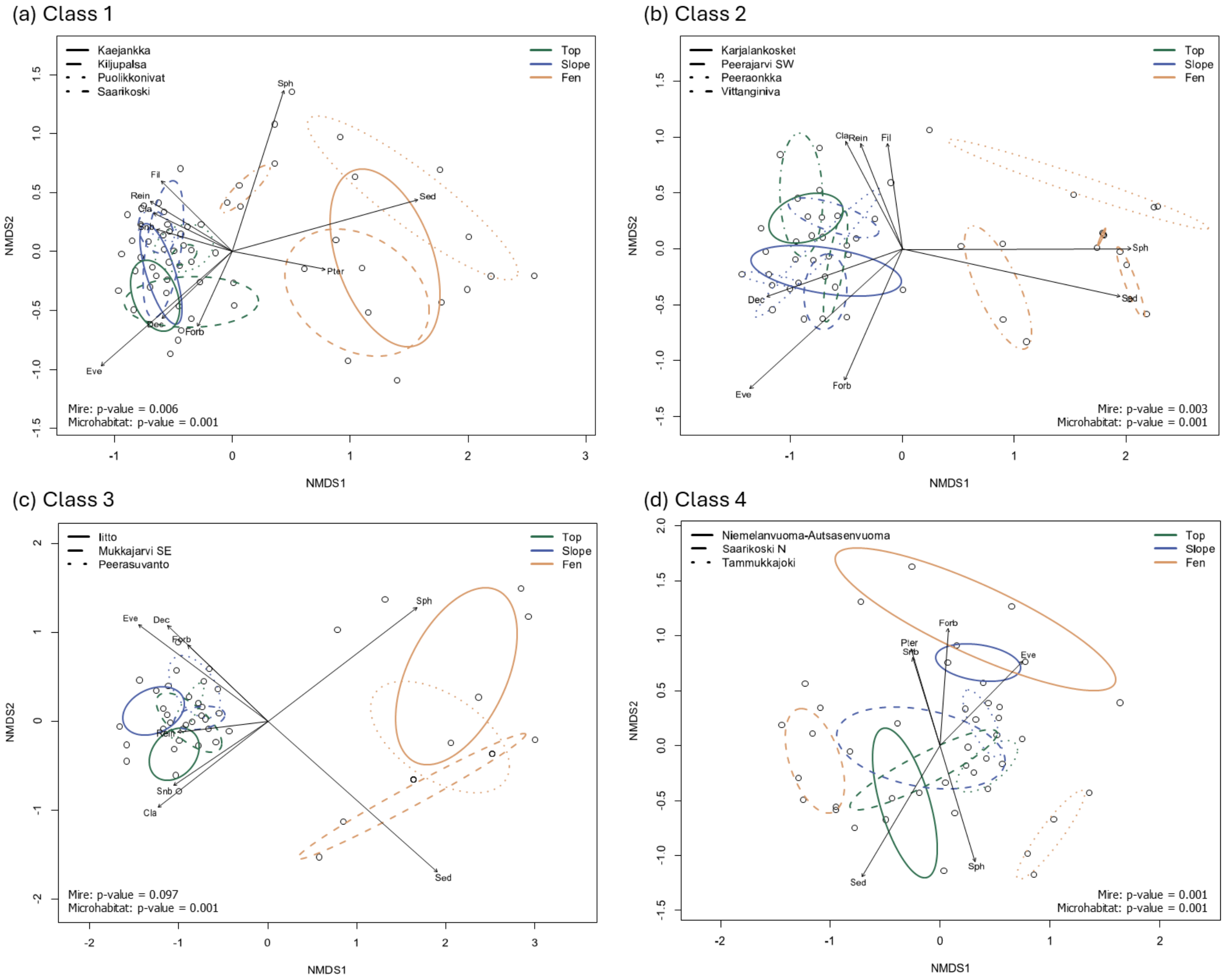


**Figure S1.** Non-metric multidimensional scaling (NMDS) illustrating the shifts in vegetation community compositions of different microhabitats in study mires across the permafrost degradation classes (Classes 1–4; a–d). Ellipses illustrate 95 % confidence intervals. The significances of study mires and microhabitats are derived from PERMANOVA. Arrows represent significant ($p < 0.05$) correlation vectors for the cover (%) of different vascular plant, bryophyte, and lichen growth forms (see Table S4 for $r^2$ and p-values). Abbreviated growth forms refer to deciduous shrubs (Dec.), evergreen shrubs (Eve.), forbs (Forb.), pteridophytes (Pter.), sedges (Sed.), Polytrichales (Pol.), bryophytes with sparsely branched or unbranched shoots (Snb.), Sphagnum (Sph.), Cladooniform fruticose (Cla.), sprawling filamentous fruticose (Fil.), and reindeer lichens (Rein.).

**Table S3.** $r^2$ and p-values (in parentheses) for the statistically significant ($p < 0.05$) correlation vectors representing the cover (%) of different vascular plant, bryophyte, and lichen growth forms used in the non-metric multidimensional scaling (NMDS) in Figure 3.

| | All habitats | Top | Slope | Fen |
|---|---|---|---|---|
| **Vascular plants** | | | | |
| Deciduous shrubs | 0.181(0.001) | 0.167(0.005) | - | - |
| Evergreen shrubs | 0.447(0.001) | 0.462(0.001) | 0.208(0.001) | 0.201(0.001) |
| Forbs | 0.178(0.001) | - | - | 0.147(0.007) |
| Grasses | - | - | 0.263(0.008) | - |
| Pteridophytes | 0.061(0.013) | - | - | - |
| Sedges | 0.716(0.001) | 0.746(0.001) | 0.288(0.002) | 0.138(0.010) |
| **Bryophytes** | | | | |
| Liverworts | 0.044(0.013) | - | - | - |
| Polytrichales | 0.039(0.022) | - | - | 0.103(0.003) |
| Shoots not or sparsely branched | 0.100(0.001) | 0.160(0.004) | - | - |
| Shoots branched | 0.039(0.041) | - | - | 0.177(0.003) |
| Sphagnum mosses | 0.382(0.001) | 0.492(0.001) | - | - |
| **Lichens** | | | | |
| Crustaceous | - | 0.164(0.010) | - | - |
| Cladooniform fruticose | 0.248(0.001) | 0.146(0.008) | 0.142(0.018) | - |
| Sprawling filamentous fruticose | 0.047(0.012) | 0.160(0.012) | - | - |
| Reindeer lichens | 0.191(0.001) | 0.182(0.003) | 0.176(0.010) | - |

**Table S4.** $r^2$ and p-values (in parentheses) for the statistically significant ($p < 0.05$) correlation vectors representing the cover (%) of different vascular plant, bryophyte, and lichen growth forms used in the non-metric multidimensional scaling (NMDS) in Figure S1.

| | Class 1 | Class 2 | Class 3 | Class 4 |
|---|---|---|---|---|
| **Vascular plants** | | | | |
| Deciduous shrubs | 0.164(0.005) | 0.272(0.001) | 0.319(0.003) | - |
| Evergreen shrubs | 0.525(0.001) | 0.518(0.001) | 0.444(0.001) | 0.243(0.003) |
| Forbs | 0.117(0.028) | 0.231(0.005) | 0.193(0.010) | 0.238(0.005) |
| Grasses | - | - | - | - |
| Pteridophytes | 0.155(0.005) | - | - | 0.177(0.005) |
| Sedges | 0.643(0.001) | 0.617(0.001) | 0.868(0.001) | 0.403(0.002) |
| **Bryophytes** | | | | |
| Liverworts | - | - | - | - |
| Polytrichales | - | - | - | - |
| Shoots not or sparsely branched | 0.108(0.029) | - | 0.221(0.003) | - |
| Shoots branched | - | - | - | 0.147(0.041) |
| Sphagnum mosses | 0.498(0.001) | 0.643(0.001) | 0.596(0.001) | 0.253(0.004) |
| **Lichens** | | | | |
| Crustaceous | - | - | - | - |
| Cladooniform fruticose | 0.133(0.018) | 0.207(0.006) | 0.329(0.001) | - |
| Sprawling filamentous fruticose | 0.174(0.010) | 0.129(0.028) | - | - |
| Reindeer lichens | 0.161(0.007) | 0.176(0.012) | 0.141(0.032) | - |

**Table S5.** Indicator species pairs for the plant communities of different microhabitats (a–c) in each permafrost degradation class. 'A' refers to specificity, describing the degree to which an indicator assemblage is restricted to a given microhabitat. 'B' refers to fidelity, which describes how consistently the indicator occurs within that microhabitat; while 'sqrtIV' is the square root of the indicator value, summarizing overall indicator strength by combining A and B. The five highest-ranking indicator species pairs (based on sqrtIV) are shown for each class–microhabitat combination; where fewer than five significant pairs were available, all are presented.

| Degradation class | Indicator assemblages | A | B | sqrtIV | p-value |
|---|---|---|---|---|---|
| **(a) Top communities** | | | | | |
| Class 1 | 1.1. *Cladonia arbuscula* + *Flavocetraria nivalis* | 0.595 | 0.850 | 0.711 | 0.005 |
| | 1.2. *Cladonia pleurota* + *F. nivalis* | 0.529 | 0.750 | 0.630 | 0.005 |
| | 1.3. *F. nivalis* + *Rubus chamaemorus* | 0.433 | 0.850 | 0.606 | 0.005 |
| | 1.4. *Andromeda polifolia* + *C. arbuscula* | 0.461 | 0.750 | 0.588 | 0.005 |
| | 1.5. *A. polifolia* + *C. pleurota* | 0.529 | 0.650 | 0.587 | 0.005 |
| Class 2 | 2.1. *Cladonia sulphurina* + *Ichmadophila ericetorum* | 0.920 | 0.250 | 0.480 | 0.005 |
| | 2.2. *C. pleurota* + *I. ericetorum* | 0.867 | 0.250 | 0.465 | 0.005 |
| | 2.3. *F. nivalis* + *Polytrichum strictum* | 0.459 | 0.438 | 0.448 | 0.015 |
| | 2.4. *A. polifolia* + *P. strictum* | 0.401 | 0.500 | 0.448 | 0.005 |
| | 2.5. *Cladonia gracilis* + *Dicranum elongatum* | 0.517 | 0.438 | 0.440 | 0.005 |
| Class 3 | 3.1. *Flavocetraria cuculata* + *Ptilidium ciliare* | 0.988 | 0.563 | 0.746 | 0.005 |
| | 3.2. *Betula nana* + *P. ciliare* | 0.556 | 0.750 | 0.646 | 0.005 |
| | 3.3. *Empetrum nigrum* + *P. ciliare* | 0.554 | 0.750 | 0.644 | 0.005 |
| | 3.4. *B. nana* + *F. cuculata* | 0.596 | 0.688 | 0.640 | 0.005 |
| | 3.5. *C. pleurota* + *P. ciliare* | 0.793 | 0.500 | 0.630 | 0.005 |
| Class 4 | 4.1. *R. chamaemorus* + *Vaccinium oxycoccos* | 0.521 | 0.500 | 0.510 | 0.005 |
| | 4.2. *Sphagnum russowii* + *V. oxycoccos* | 0.869 | 0.286 | 0.498 | 0.005 |
| | 4.3. *Eriophorum vaginatum* + *Stramiergon stramineum* | 1.000 | 0.214 | 0.463 | 0.005 |
| | 4.4. *E. vaginatum* + *Trichophorum cespitosum* | 1.000 | 0.214 | 0.463 | 0.005 |
| | 4.5. *A. polifolia* + *V. oxycoccos* | 0.428 | 0.500 | 0.463 | 0.005 |
| **(b) Slope communities** | | | | | |
| Class 1 | 1.1. *F. nivalis* + *Hepaticae* spp. | 0.802 | 0.350 | 0.530 | 0.005 |
| | 1.2. *F. nivalis* + *V. oxycoccos* | 0.914 | 0.250 | 0.478 | 0.005 |
| | 1.3. *D. elongatum* + *P. strictum* | 0.639 | 0.250 | 0.478 | 0.015 |
| | 1.4. *Hepaticae* spp. + *P. strictum* | 0.510 | 0.400 | 0.452 | 0.005 |
| | 1.5. *Cladonia rangiferina* + *P. strictum* | 0.471 | 0.400 | 0.434 | 0.005 |
| Class 2 | 2.1. *E. nigrum* + *V. oxycoccos* | 0.593 | 0.250 | 0.385 | 0.025 |
| | 2.2. *E. nigrum* + *Pleurozium schreberi* | 0.527 | 0.250 | 0.363 | 0.005 |
| | 2.3. *P. schreberi* + *Vaccinium vitis-idaea* | 0.520 | 0.250 | 0.360 | 0.010 |
| | 2.4. *B. nana* + *P. schreberi* | 0.446 | 0.250 | 0.334 | 0.010 |
| | 2.5. *P. strictum* + *Vaccinium uliginosum* | 0.415 | 0.250 | 0.322 | 0.015 |
| Class 3 | 3.1. *P. ciliare* + *R. chamaemorus* | 0.460 | 0.625 | 0.536 | 0.005 |
| | 3.2. *C. gracilis* + *P. ciliare* | 0.446 | 0.563 | 0.501 | 0.005 |
| | 3.3. *C. rangiferina* + *P. ciliare* | 0.445 | 0.500 | 0.472 | 0.015 |
| | 3.4. *C. arbuscula* + *P. ciliare* | 0.440 | 0.500 | 0.469 | 0.015 |
| | 3.5. *P. strictum* + *P. ciliare* | 0.610 | 0.313 | 0.437 | 0.020 |
| Class 4 | 4.1. *Sphagnum fallax* + *S. russowii* | 0.950 | 0.286 | 0.521 | 0.005 |
| | 4.2. *E. nigrum* + *S. fallax* | 0.836 | 0.286 | 0.489 | 0.015 |
| | 4.3. *P. schreberi* + *S. fallax* | 1.000 | 0.214 | 0.463 | 0.005 |
| | 4.4. *P schreberi* + *Vaccinium myrtillus* | 1.000 | 0.214 | 0.463 | 0.005 |
| | 4.5. *P. schreberi* + *V. oxycoccos* | 1.000 | 0.214 | 0.463 | 0.005 |
| **(c) Fen communities** | | | | | |
| Class 1 | 1.1. *Carex rostrata* + *Comarum palustre* | 0.917 | 0.250 | 0.479 | 0.005 |
| | 1.2. *C. rostrata* + *Eriophorum angustifolium* | 0.526 | 0.350 | 0.429 | 0.035 |
| | 1.3. *E. vaginatum* + *S. russowii* | 0.427 | 0.200 | 0.292 | 0.005 |
| Class 2 | 2.1. *Carex aquatilis* + *Sphagnum riparium* | 0.849 | 0.250 | 0.461 | 0.005 |
| | 2.2. *C. aquatilis* + *E. angustifolium* | 0.637 | 0.250 | 0.399 | 0.005 |
| | 2.3. *Carex canescens* + *S. riparium* | 0.507 | 0.313 | 0.398 | 0.005 |
| | 2.4. *C. canescens* + *E. angustifolium* | 0.466 | 0.313 | 0.382 | 0.005 |
| Class 3 | 3.1. *E. angustifolium* + *Sphagnum lindbergii* | 0.975 | 0.313 | 0.552 | 0.005 |
| | 3.2. *E. vaginatum* + *S. riparium* | 0.591 | 0.250 | 0.384 | 0.005 |
| Class 4 | 4.1. *C. aquatilis* + *Epilobium palustre* | 0.855 | 0.286 | 0.494 | 0.005 |
| | 4.2. *C. aquatilis* + *Hepatiecae* spp. | 1.000 | 0.214 | 0.463 | 0.005 |
| | 4.3. *C. palustre* + *S. stramineum* | 1.000 | 0.214 | 0.463 | 0.010 |

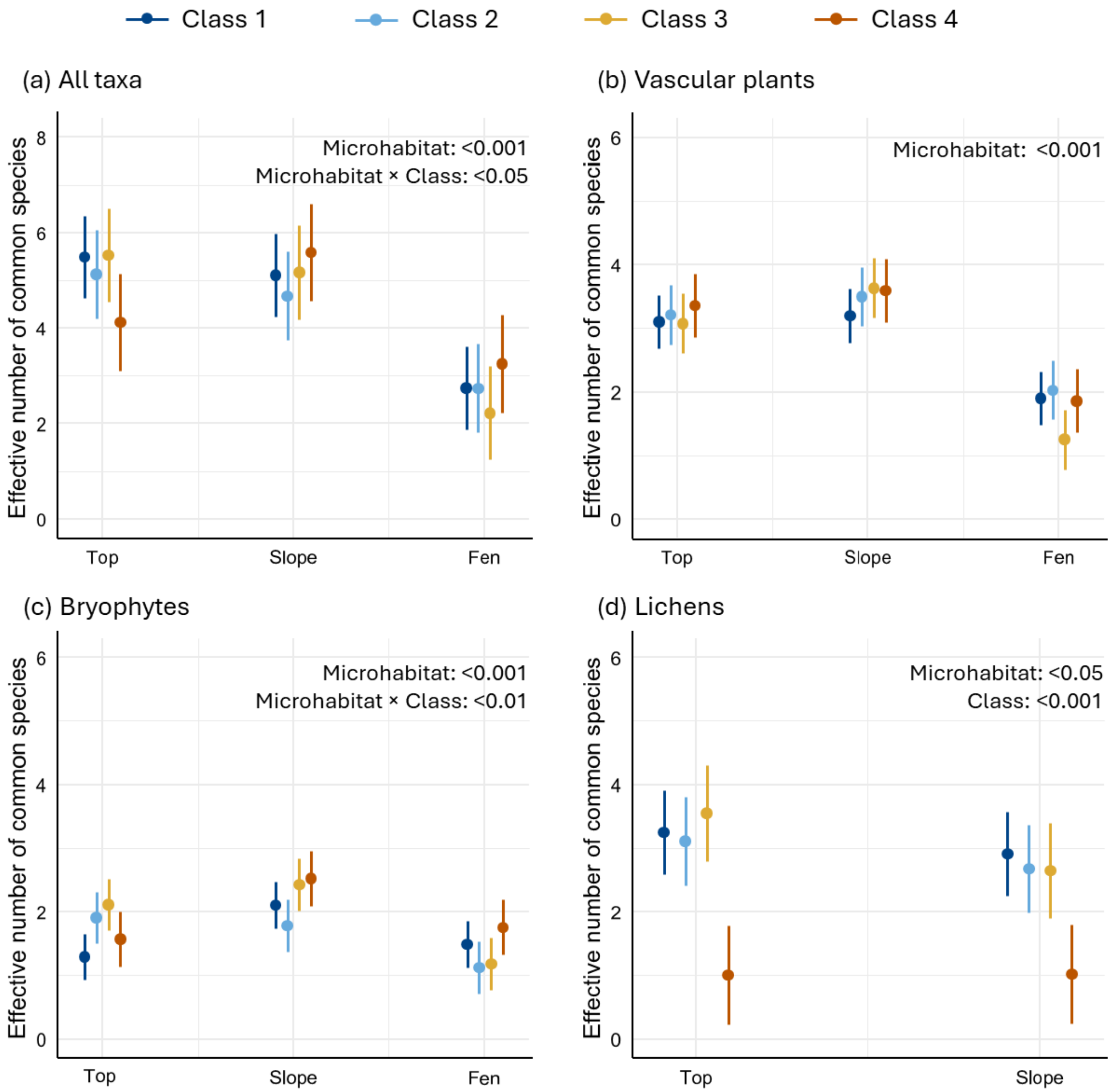


**Figure S2.** Effective number of common species (Hill number q = 1) for all taxonomic groups combined (a) and for each group separately (b–d) across the studied microhabitats. Results are based on generalized linear mixed models (GLMMs), and statistically significant explanatory variables are indicated in each panel. Note the different y-axis scale in panel (a).

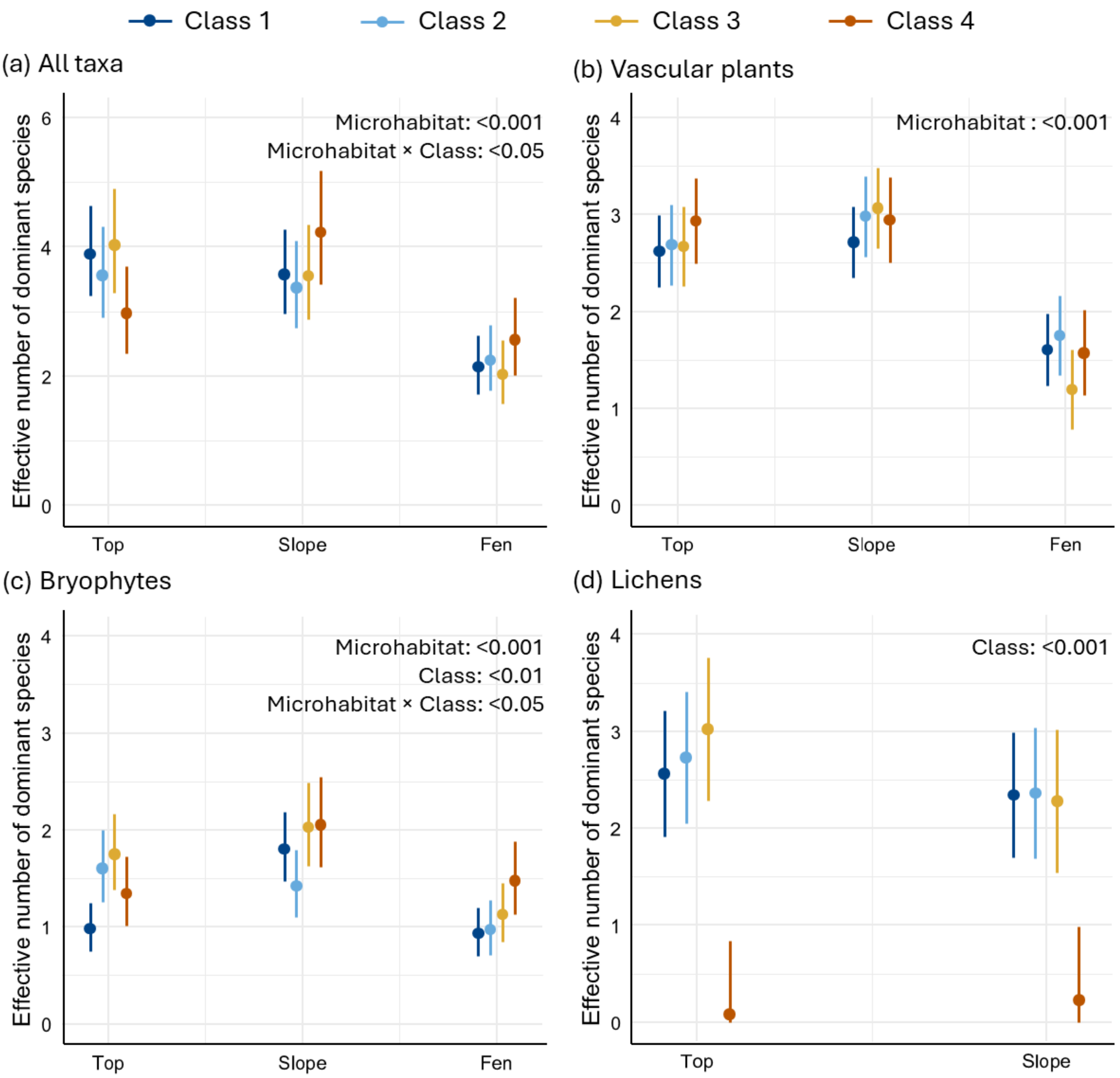


**Figure S3.** Effective number of dominant species (Hill number q = 2) for all taxonomic groups combined (a) and for each group separately (b–d) across the studied microhabitats. Results are based on generalized linear mixed models (GLMMs), and statistically significant explanatory variables are indicated in each panel. Note the different y-axis scale in panel (a).

**Table S6.** Absolute and relative covers (%) of vascular plants in different microhabitats across the permafrost degradation classes of palsa mires. Values are calculated from the plot means.

| | **Top** | | **Slope** | | **Fen** | |
|---|---|---|---|---|---|---|
| | Absolute cover | Relative cover | Absolute cover | Relative cover | Absolute cover | Relative cover |
| Class 1 | 66.65 | 69.53 | 76.65 | 73.98 | 77.35 | 61.58 |
| Class 2 | 70.75 | 66.30 | 74.19 | 63.17 | 73.44 | 48.33 |
| Class 3 | 73.81 | 71.14 | 84.63 | 72.80 | 79.06 | 49.09 |
| Class 4 | 66.29 | 42.22 | 77.36 | 60.01 | 71.79 | 50.72 |
